# Graphene Quantum Dot Oxidation Governs Noncovalent Biopolymer Adsorption

**DOI:** 10.1101/684670

**Authors:** Sanghwa Jeong, Rebecca L. Pinals, Bhushan Dharmadhikari, Hayong Song, Ankarao Kalluri, Debika Debnath, Wu Qi, Moon-Ho Ham, Prabir Patra, Markita P. Landry

**Author notes:** Corresponding authors, &. These authors contributed equally (Sanghwa Jeong, Rebecca L. Pinals).

## Abstract

The graphene quantum dot (GQD) is a carbon allotrope with a planar surface amenable for functionalization and nanoscale dimensions that confer photoluminescent properties. Collectively, these properties render GQDs an advantageous platform for nanobiotechnology applications, including as optical biosensors and delivery platforms. In particular, noncovalent functionalization offers a route to reversible modification and preservation of the pristine GQD substrate. However, a clear paradigm for GQD noncovalent functionalization has yet to be realized. Herein, we demonstrate the feasibility of noncovalent polymer adsorption to the GQD surface, with a specific focus on single-stranded DNA (ssDNA). We study how GQD oxidation level affects the propensity for polymer adsorption by synthesizing and characterizing four types of GQD substrates and investigating noncovalent polymer association to these substrates. Distinct adsorption methods are developed for successful ssDNA attachment based upon the GQD’s initial level of oxidation. ssDNA adsorption to the GQD is confirmed by atomic force microscopy, by inducing ssDNA desorption, and with molecular dynamics simulations. ssDNA is determined to adsorb strongly to no-oxidation GQDs, weakly to low-oxidation GQDs, and not at all for heavily oxidized GQDs. We hypothesize that high GQD oxygen content disrupts the graphitic carbon domains responsible for stacking with the aromatic ssDNA bases, thus preventing the formation of stable polymer-GQD complexes. Finally, we develop a more generic adsorption platform and assess how the GQD system is tunable by modifying both the polymer sequence and type.

## Introduction

Graphene is a two-dimensional hexagonal carbon lattice that possesses a host of unique properties, including exceptional electronic conductivity and mechanical strength.^1^ However, graphene is a zero-bandgap material, and this lack of bandgap limits its use in semiconducting applications.^2^ To engineer a bandgap, the lateral dimensions of graphene must be restricted to the nanoscale, resulting in spatially confined structures such as graphene quantum dots (GQDs).^3^ The bandgap of GQDs is attributed to quantum confinement,^4,5^ edge effects,^6^ and localized electron-hole pairs.^7^ Accordingly, this confers tunable fluorescence properties based upon GQD size, shape, and exogenous atomic composition. In comparison to conventional semiconductor quantum dots, GQDs are an inexpensive and less environmentally harmful alternative.^8–10^ Moreover, for biological applications, GQDs are a low toxicity, biocompatible, and photostable material that offer a large surface-to-volume ratio for bioconjugation.^8,9,11,12^

Exploiting the distinct material properties of graphene often requires or benefits from exogenous functionalization. The predominant mechanism for graphene or graphene oxide (GO) functionalization is via covalent linkage to a polymer. However, noncovalent adsorption of polymers to carbon substrates is desirable in applications requiring reversibility for solution-based manipulation and tunable ligand exchange,^13^ and preservation of the pristine atomic structure to maintain nanoscale graphene’s fluorescent characteristics.^14^ Both covalent and noncovalent functionalization of graphene and GO haver proven valuable for sensing and delivery applications. Optical sensors based on DNA-graphene or -GO hybrids have been developed for the detection of nucleic acids,^15–17^ proteins,^18,19^ small molecules,^20,21^ and metal ions.^22,23^ Additionally, modifications to GO for drug delivery applications include PEGylation for higher biocompatibility,^24,25^ covalent modification with functional groups for water solubility,^26^ noncovalent loading of anticancer drugs such as doxorubicin and camptothecin,^24,26^ and covalent linking of antibodies.^27^ Noncovalent adsorption of polymers to graphene and GO has been predicted by theory and simulations,^28,29^ and has occasionally been demonstrated experimentally.^30^ In particular, single-stranded DNA (ssDNA) of varying lengths has been experimentally shown to noncovalently attach to graphene and GO, with hydrophobic and aromatic, π-π stacking electronic interactions posited to drive assembly.^31–33^ Molecular dynamics (MD) simulations and density functional theory (DFT) modeling of these systems has enabled validation and mechanistic insight into the corresponding experimental findings.^34–36^

While noncovalent adsorption of DNA and various other polymers has been proposed by simulation and theory, and experimentally established as feasible for graphene and GO substrates, noncovalent polymer adsorption has not been fully investigated for their nanoscale counterparts: GQDs.^37^ The advantages of noncovalent GQD functionalization with biopolymers could enable two-dimensional carbon applications at the molecular scale, of relevance to study biological processes, by reducing graphene dimensions to the nanoscale. Herein, we present a facile protocol for noncovalent complexation of biopolymers to GQDs, with a focus on ssDNA. We explore the effects of GQD oxidative surface chemistry on the strength of binding interactions between surface-adsorbed polymers and GQDs, while preserving, or in some cases enabling, intrinsic GQD fluorescence. Ultimately, these results can serve as the basis for the design and optimization of polymer-GQD conjugates in various nanobiotechnology applications.

## Results and Discussion

We prepared and characterized four distinct GQD substrates of varying oxidation levels: no oxidation GQDs (no-ox-GQDs) were fabricated by coronene condensation,^38^ low oxidation GQDs (low-ox-GQDs) by intercalation-based exfoliation,^3^ medium oxidation GQDs (med-ox-GQDs) by the thermal decomposition of citric acid,^39^ and high oxidation GQDs (high-ox-GQDs) by carbon fiber cutting (Fig. 1A).^12^ X-ray photoelectron spectroscopy (XPS) was employed to quantify the differing oxidation levels among the GQD samples (Fig. 1B). The high-resolution carbon 1s (C1s) XPS signal was deconvoluted into four individual peaks attributed to sp^2^ carbon-carbon bonds (284.5 eV), hydroxyl and epoxide groups (286.1 eV), carbonyl groups (287.5 eV), and carboxyl groups (288.7 eV).^40^ The deconvoluted spectra and peak areas for each chemical bond are presented in Fig. S1. The peak area ratio of oxidized carbon (A_CO_) to sp^2^ carbon (A_CC_) decreases in the order of high-ox-GQDs (A_CO_/A_CC_=1.5) > med-ox-GQDs (0.45) > low-ox-GQDs (0.14) > no-ox-GQDs (0), as expected. Of particular note, no-ox-GQDs possessed only sp^2^ carbon, no oxidized carbon, in the C1s XPS spectrum. Atomic force microscopy (AFM) images of the GQD samples revealed the heights of high-ox-GQDs distributed between 0.5-3 nm, corresponding to 1-5 graphene layers, and the heights of med- and low-ox-GQDs between 0.5-1 nm, indicating the presence of a single graphene layer (Fig. S2). The morphology of no-ox-GQDs was separately characterized by matrix-assisted laser desorption/ionization time-of-flight mass spectroscopy (MALDI-TOF MS) due to aggregation of no-ox-GQDs during AFM sample preparation hindering equivalent AFM analysis. The single graphene layer structure of no-ox-GQDs was determined by discrete peaks in the size distribution from MALDI-TOF MS, attributed to the presence of planar dimer, trimer, tetramer, pentamer, and hexamer fused-coronene structures (Fig. S3). The fluorescence and absorbance spectra of low-, med-, and high-ox-GQDs were observed under 320 nm excitation in water (Fig. 1C and Fig. S4). The fluorescence maximum near 400 nm is described in previous literature as the intrinsic emission wavelength of GQDs with low oxidation, which is in close agreement with our own GQD samples.^10^ A red-shifted fluorescence emission peak is observed as the GQD oxidation level is increased. As previously reported, longer wavelength emission emerges due to the presence of extrinsic, defect states.^3,41^ The no-ox-GQDs were insoluble in water due to the absence of oxygen-containing functional groups, and accordingly, aggregation led to self-quenched fluorescence. Instead, the fluorescence of no-ox-GQDs was measured in hexane (Fig. 1C) and the resultant fluorescence spectrum exhibits multiple peaks originating from the size distribution of no-ox-GQD multimers. The GQD excitation-emission profiles demonstrate that the optical characteristics of low-, med-, and high-ox-GQDs depend on the excitation wavelength, where the maximum fluorescence wavelength is red-shifted as the excitation is moved to longer wavelengths. However, the fluorescence of no-ox-GQDs does not show this spectral shift (Fig. S5). This excitation-wavelength-dependence of fluorescence properties is commonly found in oxidized GQDs as a result of surface trap states introduced by functional groups and oxygen-related defects.^42^ The no-ox-GQDs do not exhibit this excitation-dependency because they do not possess any oxygen-related surface trap states. Thus, changing GQD oxidation level results in unique optical fingerprints that present an opportunity to identify GQDs by means of fluorescence profiles.

**Figure 1.**
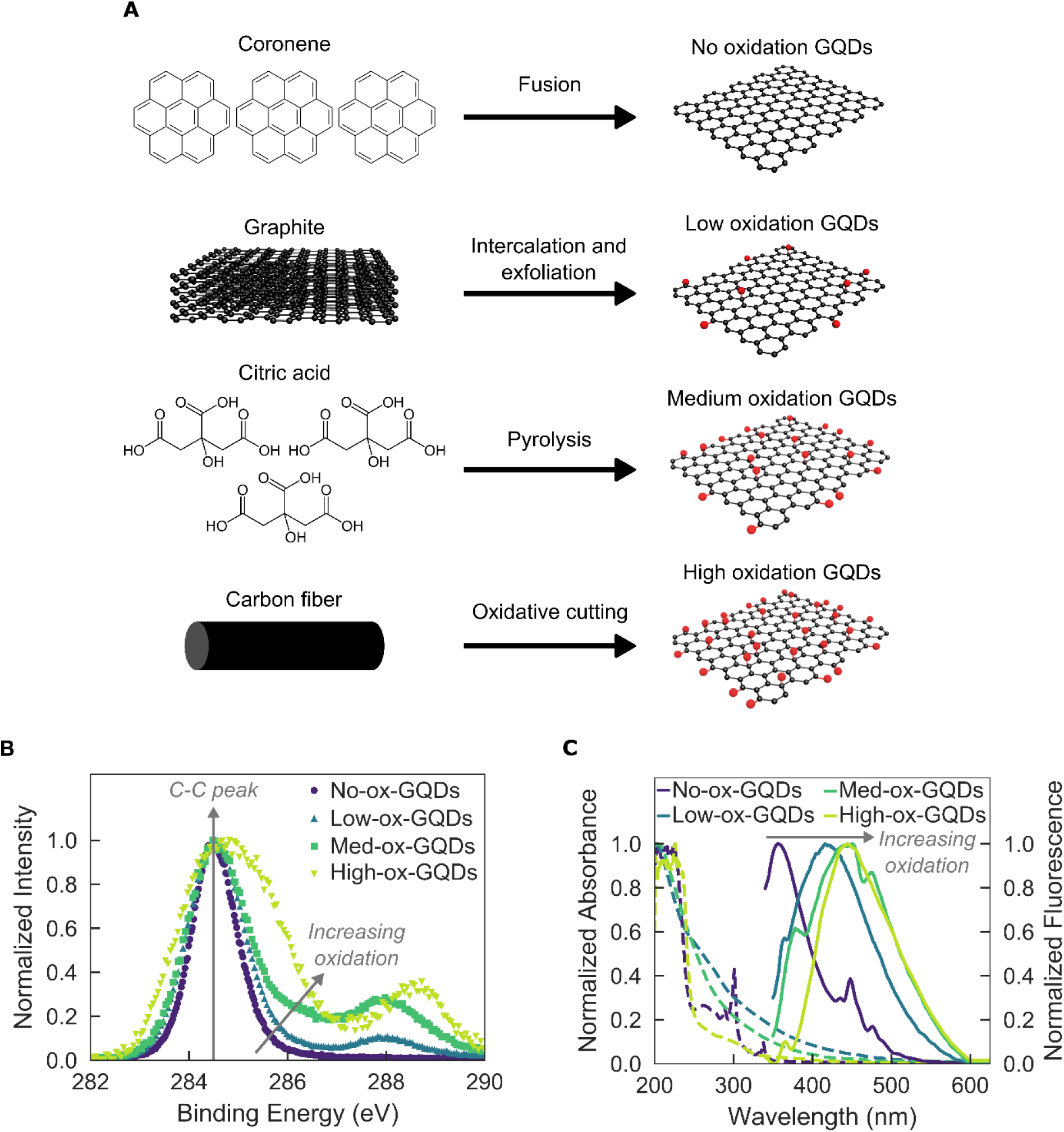
Four synthesis techniques are employed to produce graphene quantum dot (GQD) substrates of varying oxidation level. **(A)** Schematic illustration of synthesis techniques to produce no oxidation GQDs (no-ox-GQDs), low oxidation GQDs (low-ox-GQDs), medium oxidation GQDs (med-ox-GQDs), and high oxidation GQDs (high-ox-GQDs). **(B)** Normalized X-ray photoelectron spectroscopy (XPS) data of no-, low-, med-, and high-ox-GQDs. Arrows indicate the center of the C1s carbon-carbon (C-C) bond at 284.5 eV and increasing oxidation via contributions of various carbon-oxygen bonds (see Fig. S1 for peak ratios and deconvolutions). **(C)** Normalized absorbance (dashed) and fluorescence emission (solid) spectra of no-ox-GQDs in hexane solution and low-, med-, and high-ox-GQDs in water. All GQDs were excited at 320 nm.

We next studied adsorption of the ssDNA sequence (GT)_15_ onto GQDs of varying oxidation levels (Fig. 2A). This particular ssDNA oligomer was chosen for initial adsorption studies based on its known π-π stacking adsorptive properties to the surface of pristine carbon nanotubes,^43^ an analogous one-dimensional nanoscale substrate. For low-, med-, and high-ox-GQDs, ssDNA was added to GQDs in distilled (DI) water and the water was removed by vacuum evaporation to facilitate adsorption of (GT)_15_ onto the GQDs prior to GQD-ssDNA resuspension in water (mix-and-dry protocol). For no-ox-GQDs, an alternative complexation technique was employed because the as-synthesized no-ox-GQDs were insoluble in aqueous solution. Instead, the mixture of (GT)_15_ and solid no-ox-GQDs was probe-tip sonicated in PBS buffer to disperse the hydrophobic GQD aggregates and enable ssDNA adsorption. (GT)_15_ adsorption was confirmed by modulation of the intrinsic GQD fluorescence. For the initially soluble GQDs (low-, med-, and high-ox-GQDs), polymer adsorption manifests as fluorescence quenching from the original fluorescent state, whereas for the initially insoluble no-ox-GQDs, polymer adsorption manifests as fluorescence brightening from the original non-fluorescent, aggregated GQD state (Fig. 2). Fluorescence quenching was observed for low-ox-GQDs with (GT)_15_, but negligible fluorescence change was shown in the case of med- and high-ox-GQDs. These results suggest that (GT)_15_ does not adsorb to GQDs of higher oxidation levels. We confirmed fluorescence quenching of low-ox-GQDs was not a result of Förster resonance energy transfer (FRET) because there is no spectral overlap between GQD emission and ssDNA absorption, and could instead be due to charge transfer.^44^ (GT)_15_ adsorption elicits a 15 nm red-shift of the low-ox-GQD fluorescence emission peak, caused by changing polarity proximal to the GQD surface. This bathochromic shift is consistent for all biopolymers interacting with low-ox-GQDs.

**Figure 2.**
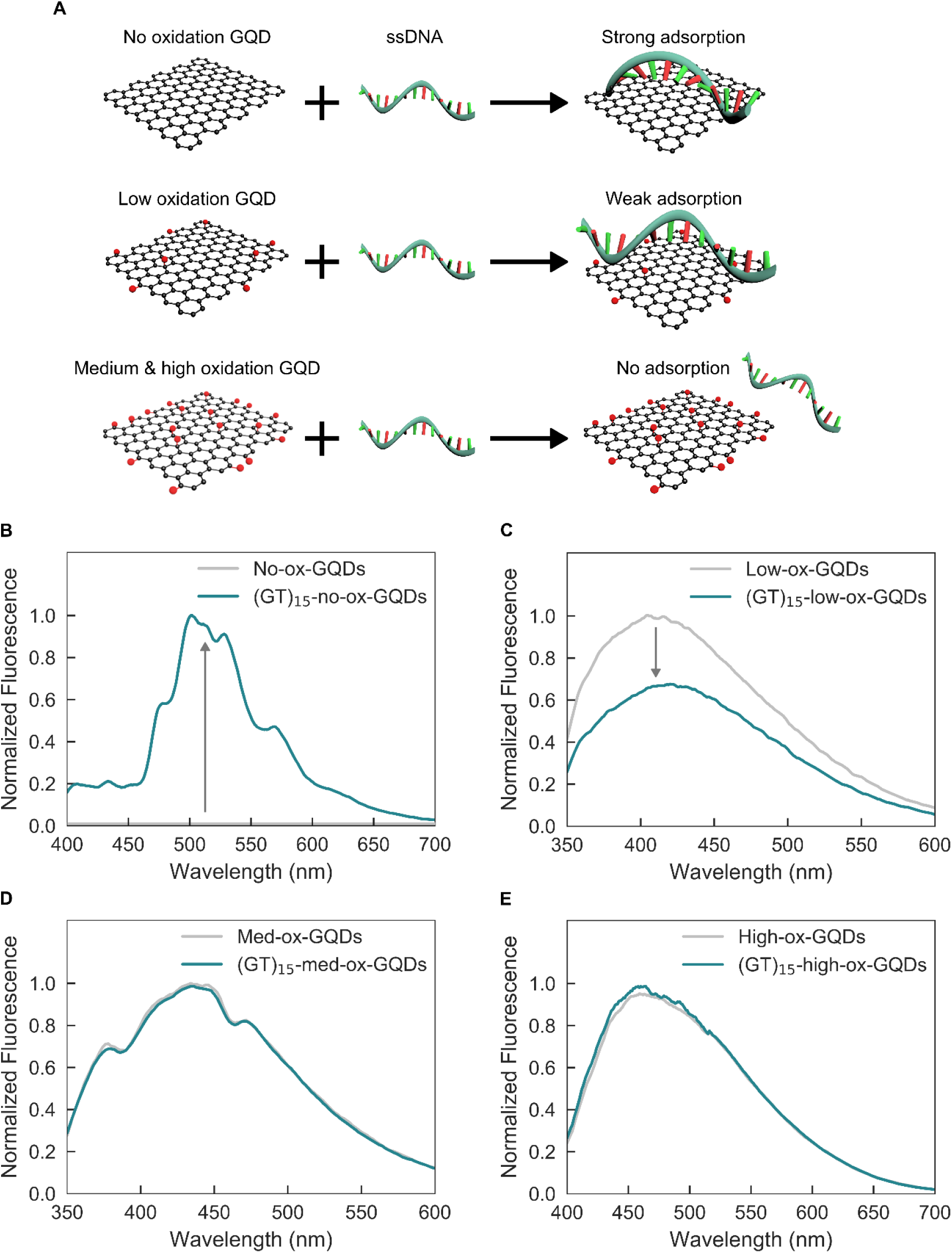
Single-stranded DNA (ssDNA)-GQD noncovalent interaction is governed by GQD oxidation level. **(A)** Schematic illustration of GQD oxidation level and resulting strength of ssDNA-GQD interaction. The noncovalent interaction between ssDNA and no-ox-GQDs is stronger than that of ssDNA and low-ox-GQDs. ssDNA does not adsorb to either med- or high-ox-GQDs. **(B-E)** Adsorption of (GT)_15_ ssDNA on the GQD surface results in GQD fluorescence modulation from before (gray) to after (blue) attempted adsorption of (GT)_15_ ssDNA for **(B)** no-ox-GQDs, **(C)** low-ox-GQDs, **(D)** med-ox-GQDs, and **(E)** high-ox-GQDs. The presence of ssDNA on the no-ox-GQDs is confirmed by an increase in fluorescence emission intensity from the initially insoluble no-ox-GQDs. The presence of ssDNA on the low-ox-GQDs results in a decrease in fluorescence intensity from the initially soluble low-ox-GQDs. No fluorescence intensity changes are observed for the med- and high-ox-GQDs, suggesting absence of ssDNA adsorption. Fluorescence spectra are normalized by the absorbance at 320 nm.

Interestingly, the simple mixing of (GT)_15_ with GQDs in the absence of drying results in only marginal fluorescence quenching for low-ox-GQDs (Fig. S6) and was accordingly ineffective in promoting ssDNA adsorption to GQDs. We hypothesize that the evaporation step is required to enable close approach of the polymers to the GQD surface by overcoming electrostatic repulsion present in solution between the negatively-charged oxidized GQDs and negatively-charged ssDNA phosphate backbone.^45^ Moreover, water molecules solvating the low-ox-GQD surface may hinder initial contact of ssDNA with low-ox-GQDs,^46^ therefore the dehydration process is necessary for ssDNA adsorption to the GQD surface. The (GT)_15_-no-ox-GQDs show a fluorescence increase from the initially non-fluorescent no-ox-GQD aggregates in aqueous solution and display the multiple absorption and emission peaks characteristic of the hexane-solubilized no-ox-GQD sample (Fig. 1C). Thus, probe-tip sonication of no-ox-GQDs with (GT)_15_ was successful in dispersing no-ox-GQDs in PBS buffer by disrupting GQD aggregates and enabling the amphiphilic ssDNA to adsorb onto the exposed hydrophobic GQD surface, thus conferring water solubility to the complex. Without ssDNA, probe-tip sonication of no-ox-GQDs in solution does not result in a stable colloidal suspension. Presence of the ssDNA on no-ox-GQDs enabled AFM analysis of the as-synthesized no-ox-GQDs and revealed heights distributed between 0.3-0.7 nm, corresponding to single graphene layer morphology (Fig. S3). The preferential adsorption of ssDNA onto no- and low-ox-GQDs implies that ssDNA adsorbs more favorably onto graphene-like carbon domains rather than oxidized carbon domains.^47^ This result points to the role of interfacial π-π electronic interactions between the GQD surface and aromatic ssDNA nitrogenous bases contributing more significantly to the overall physisorption process than hydrogen bonds between oxygen groups on the GQDs with the ssDNA. Likewise, the surface roughness created by oxygen groups on the med- and high-ox-GQDs could prevent π-π interactions of nucleobases with the graphitic surface, thus inhibiting adsorption. To the best of our knowledge, this is the first example of the noncovalent physisorption of ssDNA on GQDs, which has not been previously accomplished due to challenges arising from the small size, negative surface charge, and variable oxidation of GQD substrates.^15^ Our noncovalent attachment protocols to synthesize ssDNA-GQD complexes leads to new opportunities in developing GQD-based nucleic acid detection platforms, and further biological molecular sensing and imaging applications.

To verify the presence of ssDNA on low-ox-GQDs, we conducted AFM studies taking advantage of the well-known biotin-streptavidin interaction to impart a noticeable change in the ssDNA-GQD height profile. This assay was required because the change in height due to ssDNA adsorption alone on the GQD surface is below the limit of detection by AFM. Biotin (or vitamin H) is a small molecule with a specific and strong binding affinity for the protein streptavidin (K_d_=∼10^-14^ mol/L). Biotinylated-(GT)_15_ was adsorbed to low-ox-GQDs with the aforementioned mix-and-dry procedure to form Bio-(GT)_15_-low-ox-GQDs. Streptavidin was then mixed with the Bio-(GT)_15_-low-ox-GQDs in a 1:1 ratio of biotin: streptavidin (Bio-(GT)_15_-low-ox-GQDs + Strep) and the height profile of the resulting complexes was examined by AFM. Control samples of streptavidin mixed with non-biotinylated-(GT)_15_-low-ox-GQDs ((GT)_15_-low-ox-GQDs + Strep), biotinylated-(GT)_15_-low-ox-GQD only (Bio-(GT)_15_-low-ox-GQDs), and streptavidin only (Strep) were prepared for AFM analysis. Large biotin-streptavidin structures were frequently observed in the AFM images of Bio-(GT)_15_-low-ox-GQDs + Strep, and rarely found in the (GT)_15_-low-ox-GQDs + Strep sample, suggesting selective binding of streptavidin to the Bio-(GT)_15_-low-ox-GQDs (Fig. 3). Height distribution analysis reveals the estimated average portion of structures larger than 1.8 nm is 20.3 ± 7.3% (mean ± standard deviation) for Bio-(GT)_15_-low-ox-GQDs + Strep, compared to only 0.5 ± 0.7% for (GT)_15_-low-ox-GQDs + Strep, 0% for Bio-(GT)_15_-low-ox-GQDs, and 6.9 ± 5.0% for Strep. Here, the threshold value of 1.8 nm is the experimental sum of the GQD average height (0.6 nm) and streptavidin average height (1.2 nm). Accordingly, this confirms the formation of specific streptavidin-biotin-(GT)_15_-low-ox-GQD complexes, and thus suggests the successful noncovalent adsorption of ssDNA on the surface of low-ox-GQDs. The absence of (GT)_15_ adsorption onto med-ox-GQDs was also verified with this assay by preparing a mixture of biotinylated-(GT)_15_ and med-ox-GQDs (Bio-(GT)_15_-med-ox-GQDs) by the same method and adding streptavidin (Bio-(GT)_15_-med-ox-GQDs + Strep). Height distribution analysis reveals the estimated average portion of structures larger than 1.8 nm is 9.9 ± 0.6% for Bio-(GT)_15_-med-ox-GQDs + Strep, close to the control value of 7.5 ± 2.6% obtained for non-specific adsorption of streptavidin onto med-ox-GQDs and (GT)_15_ lacking biotin (Fig. S7). This result, in corroboration with the lack of fluorescence quenching, verifies that ssDNA does not form stable adsorbed structures with med-ox-GQDs.

**Figure 3.**
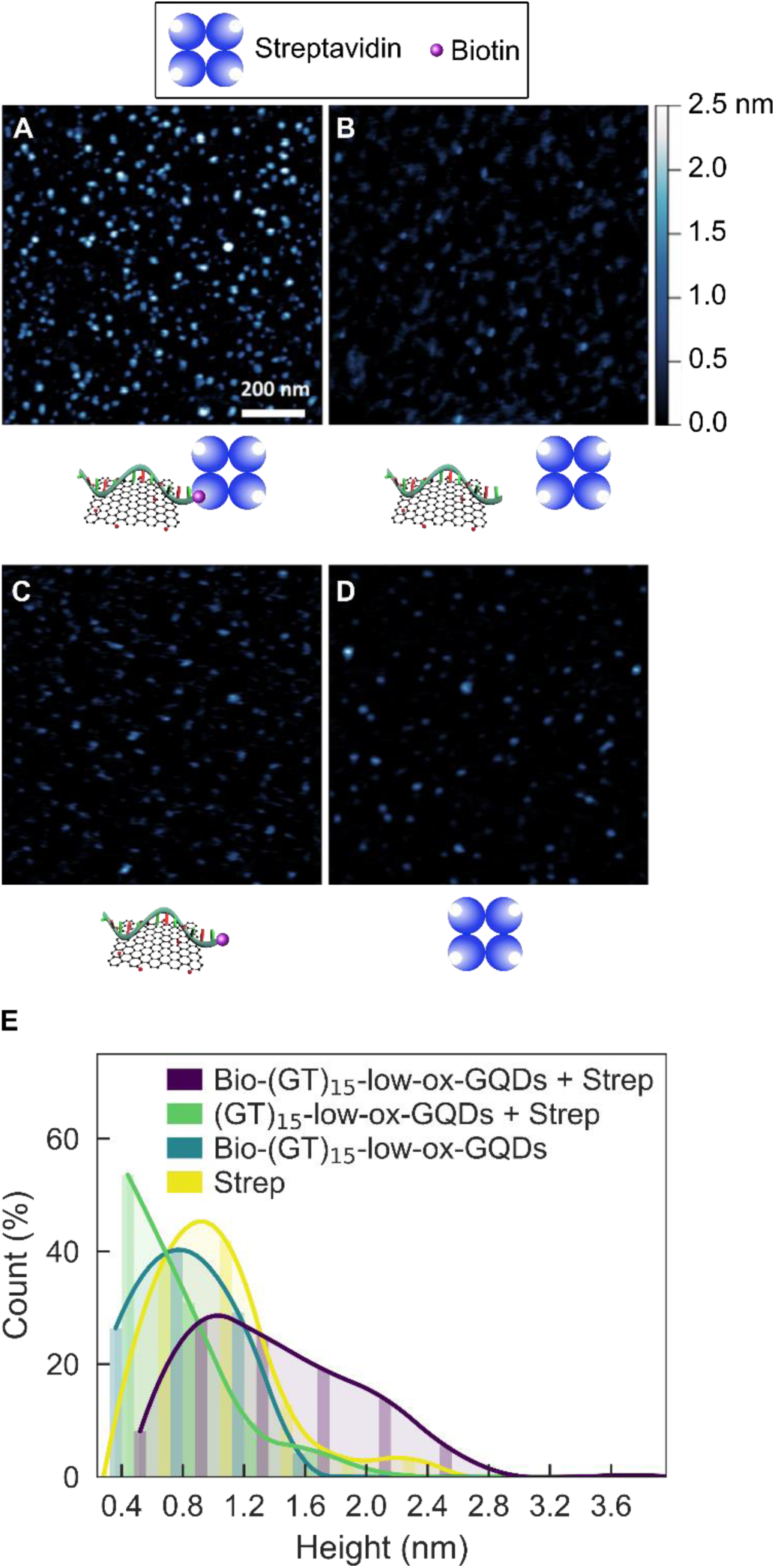
ssDNA adsorption to low-ox-GQDs is verified by Atomic Force Microscopy (AFM). AFM images and accompanying schematics for **(A)** biotinylated-(GT)_15_-low-ox-GQDs and streptavidin (Bio-(GT)_15_-low-ox-GQD + Strep), **(B)** (GT)_15_-low-ox-GQDs and streptavidin ((GT)_15_-low-ox-GQD + Strep), **(C)** biotinylated-(GT)_15_-low-ox-GQDs (Bio-(GT)_15_-low-ox-GQD), and **(D)** streptavidin (Strep). Significantly larger heights in (A) are likely due to biotin-streptavidin binding via the biotinylated-ssDNA, which is adsorbed to the low-ox-GQD surface, absent in (B), (C), and (D). **(E)** Corresponding height distribution histograms. Bin width is 0.4 nm and curve fits are added to guide the eye.

To examine the effect of ssDNA nucleotide sequence on GQD adsorption affinity, we investigated the adsorption affinities of three ssDNA oligomers of the same length but different nucleotide identities: poly-adenine, A_30_; poly-cytosine, C_30_; and poly-thymine, T_30_. Poly-guanine, G_20_, a shorter ssDNA oligomer than other ssDNA candidates, was used as the poly-G model case because of commercial unavailability of longer poly-G ssDNA oligomers. To adsorb ssDNA polymers to low-ox-GQDs, each A_30_, C_30_, G_20_ and T_30_ ssDNA oligomer was mixed and dried with low-ox-GQDs to form the ssDNA-GQD complexes: A_30_-low-ox-GQDs, C_30_-low-ox-GQDs, G_20_-low-ox-GQDs, and T_30_-low-ox-GQDs. Following ssDNA adsorption, the integrated fluorescence intensity of low-ox-GQDs decreased by 76.1 ± 8.2% (mean ± standard deviation) for A_30_, 85.1 ± 1.9% for G_20_, 72.0 ± 6.9% for T_30_ on average, and maintained the original value for C_30_ (Fig. 4A and Fig. S8). These results suggest that A_30_, G_20_, and T_30_ adsorb onto the low-ox-GQD surface, while C_30_ does not adsorb. We repeated the AFM studies with low-ox-GQD substrates to which we adsorbed biotinylated-C_30_ and mixed this construct with streptavidin (Bio-C_30_-low-ox-GQDs + Strep) to further investigate whether C_30_ adsorbs to low-ox-GQDs. As a positive control for adsorption, biotinylated-T_30_ was prepared and incubated with streptavidin (Bio-T_30_-low-ox-GQDs + Strep). AFM imaging of the biotinylated-C_30_, low-ox-GQD, and streptavidin mixture demonstrated that Bio-C_30_-low-ox-GQDs and Strep were observed as separate structures, while Bio-T_30_-low-ox-GQDs + Strep displayed heights consistent with the larger, assembled complex. Specifically, the height distribution analysis showed the estimated average portion of structures larger than 1.8 nm is 1.6 ± 0.4% for Bio-C_30_-low-ox-GQDs and Strep, which is significantly lower than the value of 12.6 ± 8.5% from the Bio-T_30_-low-ox-GQDs + Strep sample (Fig. 4B, C, D). These results, consistent with our fluorescence quenching assay, suggest that C_30_ does not adsorb onto the low-ox-GQD surface and that the binding affinity of poly-C is lower than poly-A, G and T. Previously, Sowerby *et. al*. reported the adsorption affinity of the four DNA bases onto graphite (as determined by column chromatography) in decreasing order of guanine > adenine > thymine > cytosine,^48^ in accordance with our results showing a low adsorption affinity of cytosine to GQDs. Likewise, for pyrimidine homopolymers studied with chemical force microscopy, T_50_ required a much stronger peeling force of 85.3 pN from a graphite surface as compared 60.8 pN for C_50_.^49^ Conversely, a recent study suggested that poly-C interacts with a carboxylated GO surface more strongly than poly-T or poly-A.^50^ This result was attributed to the fact that poly-C ssDNA easily forms secondary structures, enabling hydrogen bonding interactions between the folded ssDNA and GO that drive the adsorption process.^34^ However, our low-ox-GQDs contain significantly less oxidative functional groups (A_CO_/A_CC_=0.14) available for hydrogen bonding in comparison to common GO (A_CO_/A_CC_∼0.36)^51^, therefore we conclude that C_30_ was not able to interact with the same binding modalities as shown with non-nanoscale GO.

**Figure 4.**
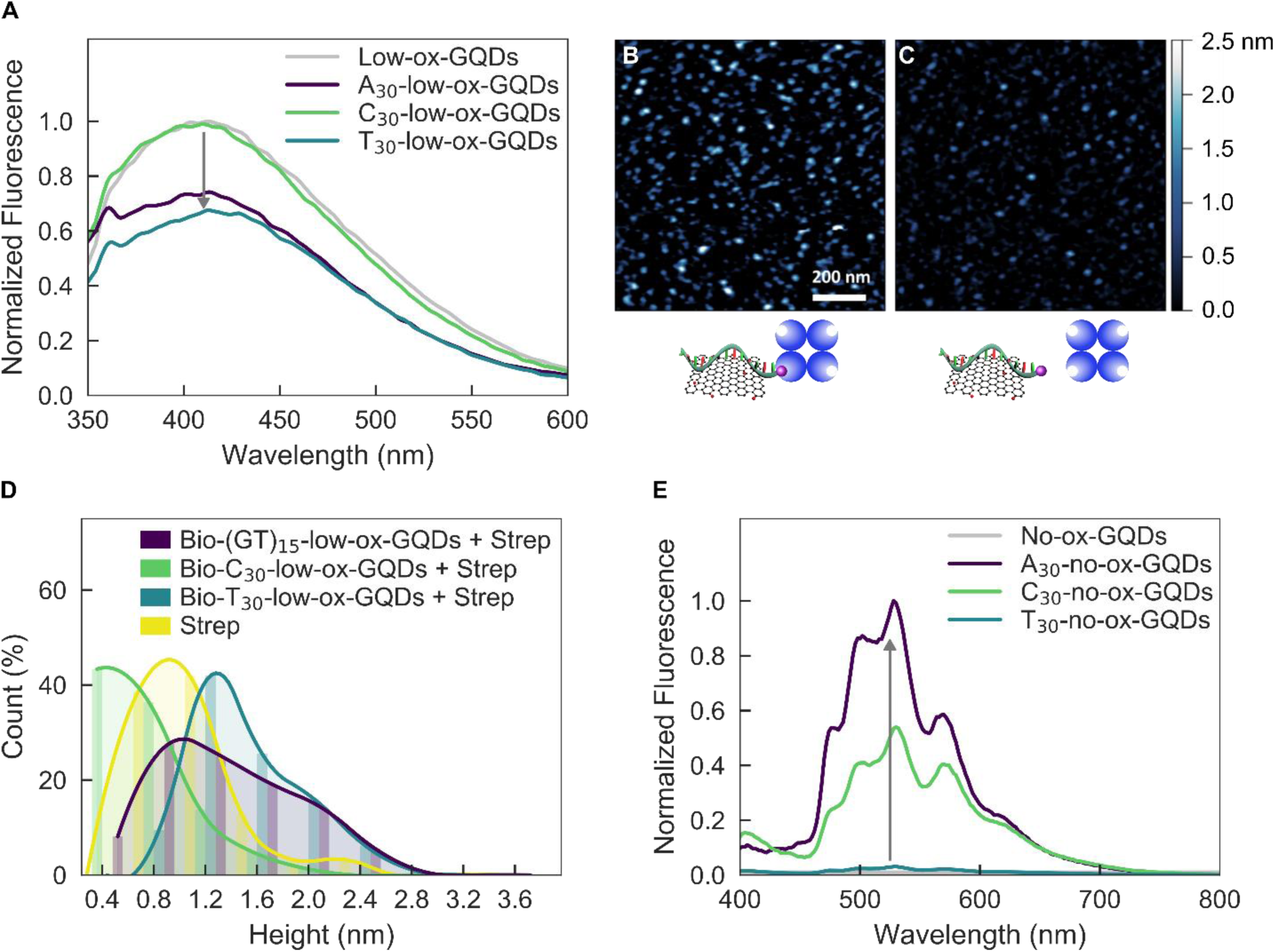
Propensity of ssDNA adsorption to low- and no-ox-GQDs depends on ssDNA sequence. **(A)** Fluorescence spectra of low-ox-GQDs (gray) and low-ox-GQDs with either A_30_ (A_30_-low-ox-GQDs), C_30_ (C_30_-low-ox-GQDs), or T_30_ (T_30_-low-ox-GQDs) adsorbed by the mix-and-dry process. AFM images and accompanying schematics for **(B)** biotinylated-T_30_-low-ox-GQDs and streptavidin (Bio-T_30_-GQD + Strep) and **(C)** biotinylated-C_30_-low-ox-GQDs and streptavidin (Bio-C_30_-GQD + Strep). **(D)** Corresponding height distribution histograms. Bin width is 0.4 nm and curve fits are added to guide the eye. **(E)** Fluorescence spectra of no-ox-GQDs (gray) and no-ox-GQDs with either A_30_ (A_30_-no-ox-GQDs), C_30_ (C_30_-no-ox-GQDs), or T_30_ (T_30_-no-ox-GQDs) adsorbed by probe-tip sonication. All GQD fluorescence spectra are normalized by the absorbance at 320 nm.

To further study the sequence dependence of ssDNA adsorption to GQD substrates, we studied the ssDNA adsorption affinity of A_30_, C_30_, and T_30_ ssDNA oligomers to no-ox-GQDs. A_30_, C_30_, and T_30_ ssDNA oligomers were probe-tip sonicated with water-insoluble no-ox-GQDs. All three ssDNA sequences resulted in stable colloidal dispersions of ssDNA-coated no-ox-GQDs. The relative fluorescence intensities normalized by the absorbance at the excitation wavelength (320 nm) establishes the fluorescence quantum yield order as A > C > T (Fig. 4D). However, it is noteworthy that this order does not directly reflect the adsorption affinity of each polynucleotide, as the fluorescence intensity is correlated to both nucleotide-specific adsorption affinity and quenching ability. The selective adsorption of C_30_ onto no-ox-GQDs, and its lack of adsorption to low-ox-GQDs, implies higher adsorption proclivity for C_30_ on no-ox-GQDs. We attribute the selective ability of C_30_ to adsorb to no-ox-GQDs over low-ox-GQDs to π-π stacking interactions for no-ox-GQDs that are strong enough to unfold the secondary structure of C_30_, but insufficient for the analogous low-ox-GQD case.

Next, we investigated ssDNA desorption from ssDNA-coated low-ox-GQDs and no-ox-GQDs by using high temperature and complementary ssDNA (cDNA) methods. To study the effect of high temperature, ssDNA-GQD samples of A_30_-, C_30_-, and T_30_ on either no- or low-ox-GQDs were prepared. The fluorescence intensity of each GQD sample was measured at room temperature before and after heating samples to 50 °C for 2 hours to attempt desorption of ssDNA from the GQD (Fig. S9A, B). As expected, no fluorescence change was observed after heating pristine low-ox-GQDs and C_30_-low-ox-GQDs because these samples did not initially have surface-adsorbed ssDNA. Fluorescence intensity of A_30_- and T_30_-low-ox-GQDs increased after heating, indicating that 47.4% of A_30_ and 30.7% of T_30_ desorbed from the low-ox-GQD surface upon heating to 50 °C. In comparison, fluorescence of all ssDNA-no-ox-GQDs maintained their initial intensity after heating to 50 °C, suggesting this heat treatment was insufficient to desorb ssDNA from the pristine no-ox-GQD carbon lattice. When ssDNA-no-ox-GQDs were instead heated to 95 °C for 2 hours, fluorescence intensities of all groups significantly decreased, indicating that 41.9% of A_30_, 43.6% of C_30_, and 39.3% of T_30_ desorbed from the no-ox-GQD surface, suggesting this higher heat treatment is required to induce ssDNA desorption and subsequent aggregation of insoluble no-ox-GQDs (Fig. S10). This difference of temperature stability implies the adsorption affinity of ssDNA on the GQD surface is much stronger for no-ox-GQDs than for low-ox-GQDs. A recent MD simulation study reported that the estimated binding free energy between T_20_ ssDNA and GO increased significantly when the oxygen content of GO is reduced to below 10%.^34^ Accordingly, we hypothesize that the GQD oxidation level is directly related to the adsorption affinity between ssDNA and GQDs. The stronger adsorption affinity of ssDNA onto the no-ox-GQDs results from the increased sp^2^ graphitic carbon content available for π-π stacking interaction with ssDNA and reduced GQD negative surface charge for electrostatic repulsion. The adsorption stability of ssDNA on GQDs was also studied with a hybridization assay, where ssDNA complementary to the adsorbed sequence, cDNA, hybridizes in solution phase with the GQD surface-adsorbed ssDNA. It is known that double-stranded DNA has a low adsorption affinity for GO surfaces, and this property has been previously used to study the adsorption affinity of ssDNA by cDNA-induced desorption.^47^ The cDNA oligomer (AC)_15_ was added to either (GT)_15_-low-ox-GQD or (GT)_15_-no-ox-GQD solutions in five-fold excess relative to (GT)_15_. The resulting fluorescence profile of each GQD solution was measured 2 hours following addition of (AC)_15_ and compared with the initial fluorescence profile (Fig. S9C, D). Fluorescence of low-ox-GQDs decreased by 68% upon initial (GT)_15_ adsorption, and was recovered to 88% of the initial low-ox-GQD fluorescence due to ssDNA desorption in the presence of cDNA. On the other hand, fluorescence of (GT)_15_-no-ox-GQDs maintained the initial fluorescence value after adding cDNA. These results further substantiate our conclusion that ssDNA adsorbs to no-ox-GQDs more strongly than to low-ox-GQDs.

To understand the time-dependent energetics and structures of the ssDNA-GQD binding process, we performed MD simulations of polynucleotide ssDNA oligomers adsorbing to GQDs of varying oxidation levels. To investigate how GQD surface polarity affects ssDNA adsorption, we analyzed non-bonding interaction energies between A_30_ ssDNA and differentially oxidized GQDs during a 100 ns MD simulation. We utilized three types of GQDs, with 0%, 2.28%, and 17.36% oxidation (denoted as GQD-0%, GQD-2%, and GQD-17%, respectively), calculated as the ratio of oxidized carbon to sp^2^ carbon. Overall, our results indicate that ssDNA physisorption is driven by a combination of van der Waals’s (vdW) interactions and hydrogen bonding (H-bonding) to the GQD. Based on the contact area of ssDNA on the GQD surface, center-of-mass distance, and number of atoms within 5 Å of the GQD surface, A_30_ ssDNA is more closely adsorbed on less oxidized GQD surfaces, such as GQD-0% and GQD-2%, as compared to the more highly oxidized surface of GQD-17% (Fig. 5A and Fig. S11A, B). These results indicate that vdW interactions are the sole contributor towards the adsorption of A_30_ on GQD-0%, whereas H-bonding marginally contributes to the adsorption of A_30_ on GQD-2% and GQD-17% in addition to dominant vdW interactions (Fig. 5B, Fig. S11C). This result further supports the more significant role of π-π electronic interactions in comparison to hydrogen bonds between the ssDNA and oxygen groups on the GQDs. In the simulation for GQD-17%, A_30_ showed negligible contact with the GQD until 70 ns, in comparison with A_30_ contact within 20 ns for the less oxidized GQD cases. After 70 ns, transient contact of A_30_ with GQD-17% was observed, as signified by the fluctuating center-of-mass distances, the latter suggesting highly unstable physisorption of A_30_ on GQD-17%. These MD results suggest more stable adsorption of A_30_ onto less oxidized GQDs (GQD-0% and GQD-2%) as compared to GQD-17%, and agree with experimentally determined selective adsorption of ssDNA on no- and low-ox-GQDs, which is not observed in the case of med- and high-ox-GQDs.

**Figure 5.**
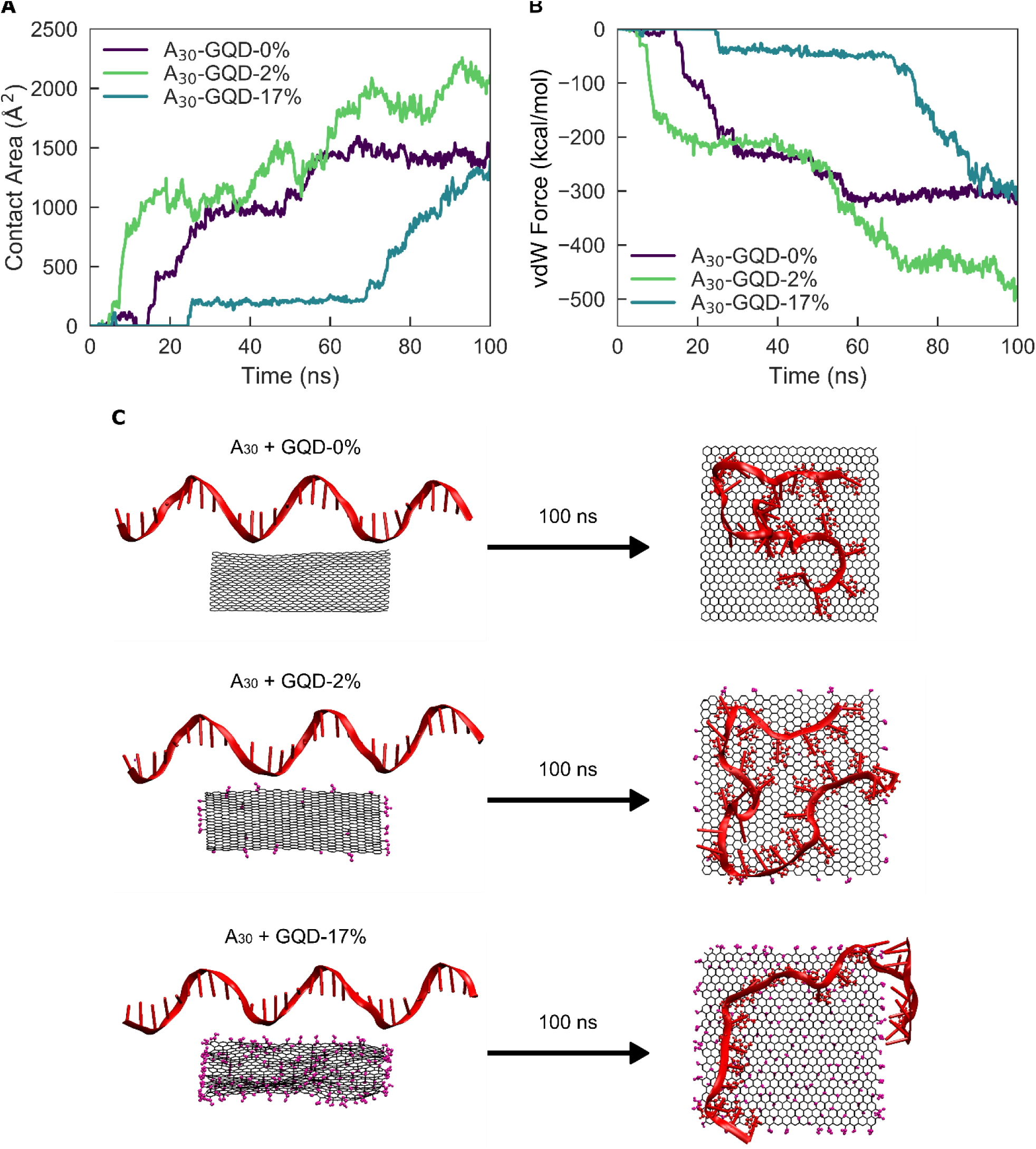
Molecular dynamics simulations confirm A_30_ ssDNA adsorption dependency on GQD oxidation level. Time-dependent **(A)** contact area and **(B)** van der Waals interactions for A_30_ ssDNA adsorbing to GQD-0%, GQD-2%, and GQD-17%. **(C)** Initial (left) and final (right) configurations of A_30_ ssDNA with GQD-0%, GQD-2%, and GQD-17% for a 100 ns simulation.

We next investigated the dependency of ssDNA-GQD adsorption on nucleotide sequence by performing MD simulations of A_30_, C_30_, and T_30_ ssDNA onto GQD-0% and GQD-2% (Fig. S12 and Fig. S13). While A_30_ and T_30_ displayed similar adsorption dynamics onto GQD-0% and GQD-2%, C_30_ adsorbed more weakly onto GQD-0% and GQD-2%, in alignment with previous studies regarding the interaction of homopolynucleotides with graphite.^48,49^ For the GQD-0% case, the simulation shows that only the 5’ end of C_30_ interacts with the GQD-0% surface, while the other end attempts self-hybridization and consequently unfolds. Interestingly, C_30_ does not show any attractive interaction with GQD-2%. This simulation result corroborates experimental results that C_30_ does not quench low-ox-GQD fluorescence and was not found in appreciable quantities on the low-ox-GQD surface by AFM. Likewise, this result supports our prior conclusion that π-π stacking interactions between the aromatic portions of ssDNA and pristine graphitic areas of GQDs can somewhat overcome the intramolecular forces holding together the C_30_ secondary structure, resulting in some adsorption of C_30_ to GQD-0% and no adsorption to GQD-2%. Overall, the MD simulations recapitulate experimental findings that GQD oxidation level determines the ssDNA interaction with and conformation on the GQD surface, and that the ssDNA-GQD-2% interaction is strongly dependent on the nucleotide sequence.

Finally, we demonstrate that this noncovalent adsorption platform is extensible to other biomolecules on GQDs. We hypothesized that planar sheet- or bilayer-forming molecules would be amenable for adsorption onto a two-dimensional GQD substrate. Accordingly, we attempted and successfully created biopolymer-GQD constructs with two such structure-forming biomolecules, phospholipids and peptoids. The phospholipid, 1,2-distearoyl-sn-glycero-3-phosphoethanolamine-N-diethylenetriaminepentaacetic acid (14:0 PE-DTPA), was adsorbed onto low-ox-GQDs with the same mix-and-dry protocol as ssDNA, and resulted in the expected fluorescence quenching of low-ox-GQDs that confirms adsorption (Fig. 6A). A peptoid is a synthetic peptide in which the variable group is attached to the amine rather than the alpha carbon, resulting in a loss of the chiral center.^52^ In particular, 36-mer peptoids with alternating ionic and hydrophobic sidechains have been designed as an amphiphilic, sheet-forming peptoid.^53^ Two peptoid sequences were tested: Peptoid-1 is a diblock of alternating *N*-(2-aminoethyl) glycine (Nae) and N-(2-phenethyl) glycine (Npe) units, abbreviated (Nae–Npe)_18_, and *N*-(2-carboxyethyl) glycine (Nce) and Npe, abbreviated (Nce–Npe)_18_. Electrostatic interactions between the amine and carboxyl groups drive solution-phase self-assembly of these 36-mers into a nanosheet morphology. Peptoid-2 is simply (Nce–Npe)_36_, with only the carboxyl sidechain present. Therefore, no amine-carboxyl ionic interactions are available to initiate assembly and this peptoid is incapable of forming stable nanosheets by itself. Probe-tip sonication of no-ox-GQDs with Peptoid-1, (Nae–Npe)_18_-(Nce–Npe)_18_, resulted in a stable Peptoid-1-no-ox-GQD suspension (Fig. 6B). The phenyl sidechains are posited to π-π stack with the no-ox-GQD basal plane in the same manner as ssDNA, thus resulting in stable constructs. Peptoid-2, (Nce–Npe)_36_, was not able to suspend the no-ox-GQDs, most likely due to the absence of electrostatic interactions between the peptoids required to form stable sheet nanostructures and presumably also a GQD surface coating. Successful synthesis of the Peptoid-1-GQD construct motivates future developments in biopolymer-GQD-based detection platforms with peptoid-mediated protein recognition.^55^ The noncovalent adsorption of biopolymers beyond ssDNA to GQDs provides a route to tune the GQD system not only by choice of GQD color and oxidation level, but additionally by polymer sequence and type.

**Figure 6.**
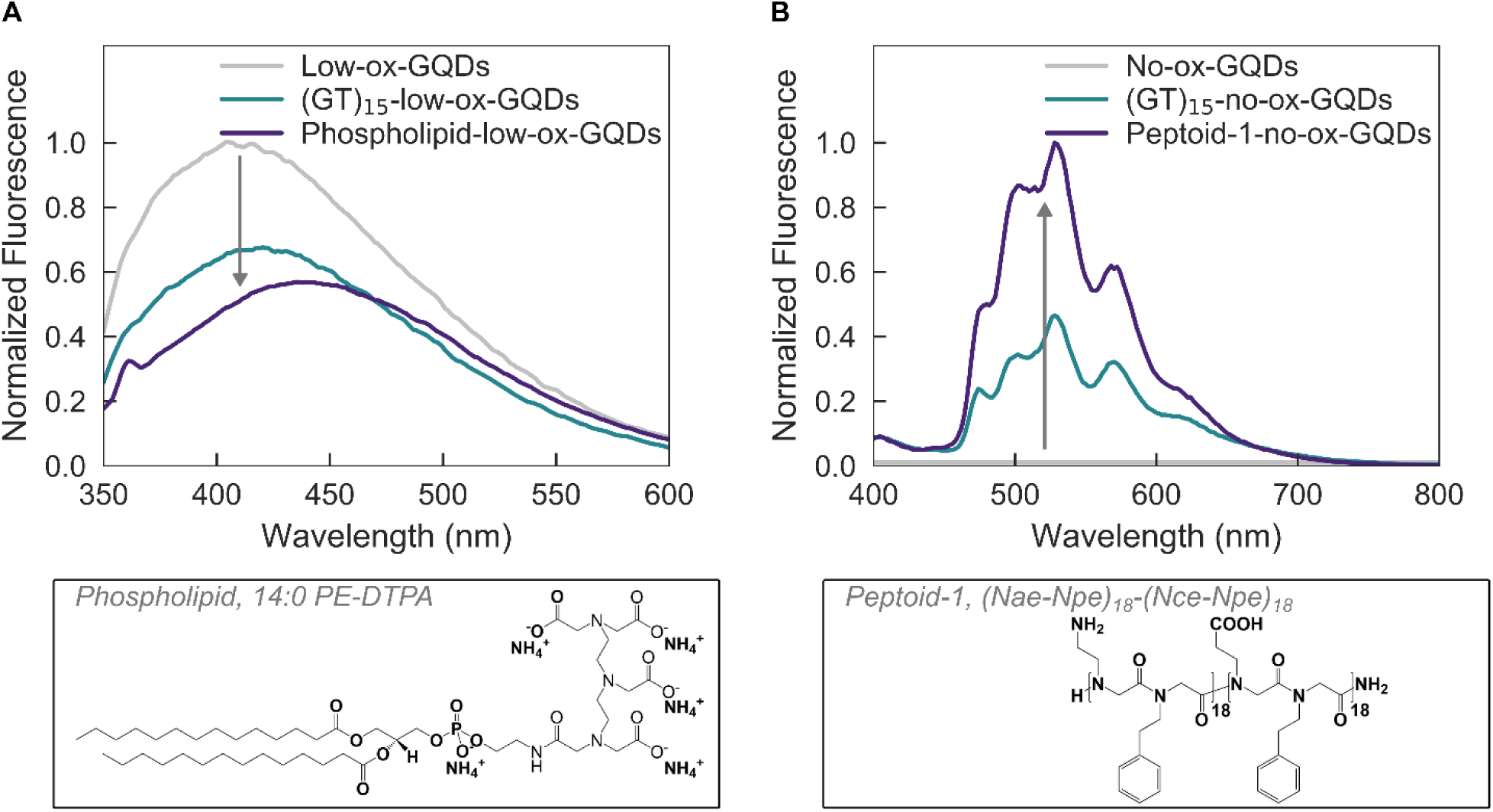
Noncovalent surface adsorption of biopolymers to low- and no-ox-GQDs is demonstrated by fluorescence modulation upon adsorption of phospholipid (14:0 PE-DTPA) and Peptoid-1 ((Nae–Npe)_18_-(Nce–Npe)_18_), respectively. **(A)** Normalized fluorescence emission spectra of low-ox-GQDs taken before (gray) and after (purple) the mix-and-dry process with phospholipid, 14:0 PE-DTPA. **(B)** Normalized fluorescence emission spectra of no-ox-GQDs taken before (gray) and after (purple) probe-tip sonication with Peptoid-1, (Nae-Npe)_18_-(Nce-Npe)_18_. The (GT)_15_-no-ox-GQDs spectrum (blue) is included for comparison. All GQD fluorescence spectra are normalized by the absorbance at 320 nm.

## Conclusions

We have demonstrated the feasibility of, and developed procedures for, noncovalent adsorption of ssDNA, phospholipids, and peptoid polymers to GQDs, confirmed the perturbative role of GQD oxidation on ssDNA adsorption, and further investigated the varying adsorption and desorption properties of ssDNA based on the GQD oxidation level and ssDNA sequence. To this end, four types of photoluminescent GQDs with different oxidation levels were fabricated. The GQD oxygen content determines the ssDNA adsorption affinity, where ssDNA can adsorb on no- and low-ox-GQD surfaces but not on med-nor high-ox-GQDs. ssDNA is adsorbed more strongly onto the no-ox-GQDs than the low-ox-GQDs, presumably because of the increased π-π stacking interactions with ssDNA and reduced electrostatic repulsion in the absence of oxygen groups. Adsorption of ssDNA on low-ox-GQDs is confirmed by fluorescence quenching and AFM studies, and the adsorption affinity of ssDNA to GQDs can be perturbed by high temperature and by hybridization with cDNA. The degree of ssDNA attachment on the low-ox-GQD surface is found to be sequence-dependent: poly-A, G, T do adsorb to low-ox-GQDs, while poly-C does not adsorb. Adsorption of ssDNA on no-ox-GQDs is confirmed by producing stable colloidal suspensions with fluorescence emission. ssDNA displays high adsorption affinity on no-ox-GQDs, which resists disruption by high temperature and cDNA. MD simulations are implemented to provide a more mechanistic understanding of the ssDNA-GQD interaction process, and these simulation results are found to corroborate experimental findings. Furthermore, we show the adsorption protocols developed for ssDNA are generic to adsorb other biopolymers, such as phospholipids and peptoids, to GQDs.

Applications of graphene-based nanomaterials are vast, and a better understanding of parameters that affect adsorption of polymers to GQDs are needed to enable future applications in diagnostics, biomolecule delivery, and sensing. We have established a method for ssDNA physisorption onto photoluminescent GQDs and studied the fundamental relation between polymer adsorption affinity and GQD oxidation level. The platform developed here can be leveraged to expand the current possibilities of designing and applying GQDs for graphene-based nanotechnologies.

## Methods

### Preparation of no oxidation GQDs (no-ox-GQDs)

No-ox-GQDs were prepared according to previous literature.^38^ Briefly, 20 mg of coronene (95%, Acros) was vacuum-sealed in a glass ampule and annealed at 500 °C for 20 hours. After cooling to room temperature, the product was loaded into a quartz tube and annealed at 500 °C for 30 min under H_2_ and Ar atmosphere (10 and 200 sccm, respectively) to remove unreacted coronene.

### Preparation of low oxidation GQDs (low-ox-GQDs)

Low-ox-GQDs were prepared by an intercalation-based exfoliation method.^3^ 20 mg of graphite powder (natural, briquetting grade, - 100 mesh, 99.9995%, UCP-1 grade, Ultra “F” purity, Alfa Aesar) and 300 mg of potassium sodium tartrate tetrahydrate (>99%, Sigma-Aldrich) were ground in a mortar and pestle under ambient conditions. The powder was transferred into a glass tube and heated in a tube furnace at 250 °C for 24 hours under Ar gas. The product powder was dispersed in 30 mL of deionized (DI) water and ultrasonicated for 10 min. The translucent, brown solution was centrifuged at 3220 g for 30 min and the supernatant solution was collected. For desalting and size selection, the solution was spin-filtered using a 100 kDa molecular weight cutoff (MWCO) centrifugal filter (Amicon Ultra-15, Ultracel, Millipore) at 3220 g for 30 min and the eluent solution was collected. The final product solution was spin-filtered with a 3 kDa centrifugal filter at 3220 g for 30 min to remove residual salts, repeated at least six times, and the remnant solution was collected.

### Preparation of medium oxidation GQDs (med-ox-GQDs)

Med-ox-GQDs were prepared according to previous literature.^39^ 2 g of citric acid (>99.5%, ACS reagent, Sigma-Aldrich) was added to a 20 mL vial and heated to 200 °C in a heating mantle. The temperature of the heating mantle was maintained for 30 min at 200 °C until citric acid was liquified into an orange solution. The solution was cooled to room temperature and added dropwise into 100 mL of 10 mg/mL NaOH solution while stirring. The pH of the med-ox-GQDs solution was neutralized to pH 7 by adding NaOH. The final product solution was spin-filtered with a 3 kDa centrifugal filter at 3220 g for 30 min to remove residual salts, repeated at least six times, and the remnant solution was collected.

### Preparation of high oxidation GQDs (high-ox-GQDs)

High-ox-GQDs were prepared according to previous literature.^12^ Briefly, 0.3 g of carbon fibers (>95%, carbon fiber veil, Fibre Glast) was added into a mixture of concentrated H_2_SO_4_ (60 mL) and HNO_3_ (20 mL). The mixture was sonicated for two hours and stirred for 24 hours at 120 °C. The solution was cooled to room temperature and diluted with DI water (800 mL). The pH of the high-ox-GQDs solution was adjusted to pH 8 by adding Na_2_CO_3_. The final product solution was spin-filtered with a 3 kDa centrifugal filter at 3220 g for 30 min to remove residual salts, repeated at least six times, and the remnant solution was collected.

### GQD characterization

XPS spectra were collected with a PHI 5600/ESCA system equipped with a monochromatic Al K_α_ radiation source (hν = 1486.6 eV). High-resolution XPS spectra were deconvoluted with MultiPak software (Physical Electronics) by centering the C-C peak to 284.5 eV, constraining peak centers to ±0.1 eV the peak positions reported in previous literature,^40^ constraining full width at half maxima (FWHM) ≤1.5 eV, and applying Gaussian-Lorentzian curve fits with the Shirley background. AFM images were collected with an MFP-3D system (Asylum) in tapping mode with an NCL-20 AFM tip (force constant = 48 N/m, Nanoworld). Optical properties of the GQDs were studied with absorbance spectroscopy (UV-3600 Plus, SHIMADZU), photoluminescence spectroscopy (Quantamaster Master 4 Lformat, Photon Technologies International), and excitation-emission profiles (Cary Eclipse, Varian). MALDI-TOF mass spectra were acquired on an Autoflex Max (Bruker) with a 355-nm laser, in the positive reflectron mode. Samples were added to CHCA matrix.

### Fabrication of ssDNA-GQD complex by mix-and-dry process

10 µg of GQDs were mixed with 10 nmol of 30-mer ssDNA (e.g. (GT)_15_) dissolved in 0.2 mL DI water. The mixture was dried for 4 hours in a 1.5 mL microcentrifuge tube under moderate vacuum (∼5 torr). The dried solid was re-dispersed in 1 mL DI water.

### Fabrication of ssDNA-no-ox-GQD complex by probe-tip sonication process

1 mg of no-ox-GQDs and 100 nmol of 30-mer ssDNA was dispersed in 1 mL PBS buffer at pH 7.4. The mixture was bath-sonicated for 2 min (Branson Ultrasonic 1800) and probe-tip sonicated for 30 min at 5 W power in an ice bath (Cole Parmer Ultrasonic Processor, 3 mm tip diameter). The product was centrifuged at 3300 g (Eppendorf 5418) for 10 minutes to remove unsuspended no-ox-GQDs and the supernatant was collected. The suspension was centrifuged at 16000 g for 1 hour to remove free ssDNA and the precipitate was collected. This purification step was repeated three times until no ssDNA was observed in the supernatant solution by absorption spectroscopy.

### Verification of ssDNA-GQD complexes by AFM

Biotinylated-(GT)_15_-low-ox-GQDs (Bio-(GT)_15_-low-ox-GQDs) were prepared by the mix-and-dry process using 5’-biotinylated-(GT)_15_ and low-ox-GQDs. To form the biotin-streptavidin complex, Bio-(GT)_15_-low-ox-GQDs, containing 10 pmol of biotinylated-(GT)_15_, and 10 pmol of streptavidin were mixed in 0.02 mL DI water. The 50-fold diluted solution was drop cast onto a mica substrate and dried by N_2_ flow. As a negative control, (GT)_15_-low-ox-GQDs were also mixed with streptavidin in the same way. AFM analysis was performed in tapping mode and the height of the GQD complex was determined as the maximum height at the GQD region in the AFM image. The average and standard deviation of the relative portion of structures larger than 1.8 nm was calculated from the height count data of multiple AFM images.

### Molecular dynamics (MD) simulations

MD simulations of ssDNA adsorption on GQDs were performed by NAMD^55,56^ using CHARMM27 and CHARMM36^57^ force field parameters for 100 ns. The obtained trajectories were visualized and analyzed using VMD^58^. The crystallographic data coordinates of A_30_, C_30_, and T_30_ ssDNA in the form of pdb files were generated using 3-D DART software^59^. The 5 nm x 5 nm GQDs with sp^2^ hybridized carbon atoms were generated using the VMD plug-in, “nanotube-builder”. The hydroxyl, carbonyl, and carboxyl groups were placed randomly on the surface and edges of the GQDs with VEGA ZZ software.^60^ The minimum distance between the ssDNA and the GQDs was 1.4 nm to maintain several ordered water layers that reduced the effects by initial status. The ssDNA and GQDs were then solvated using TIP3P water model^61^ with 150 mM sodium and chloride ions. The size of the water box was 130 × 80 × 60 Å^3^. Initial position and orientation of ssDNA were the same in all simulations.

All computations were performed at a constant temperature of 300 K and a constant pressure of 1 atm. Lennard–Jones potential parameters were set to study the cross-interaction between non-bonded atoms of ssDNA-GQD, GQD-water and ssDNA-water. All atoms, including hydrogen, were defined explicitly in all simulations. The CHARMM force field parameter files were specified to control the interaction between non-bonded atoms of a ssDNA-GQD, GQD-water and ssDNA-water. The exclude parameter was set to scaled1-4, such that all atom pairs directly connected via a linear bond and bonded to a common third atom along with all pairs connected by a set of two bonds were excluded. The electrostatic interactions for the above pairs were modified by the constant factor defined by 1-4scaling, set to 1. The cutoff distance and switching distance function were set to 14 and 10, respectively, and the switching parameter set to on, such that the van der Waals energy was smoothly truncated at the cutoff distance starting at the switching distance specified. The pair list distance (pairlistdist) was set to 14 to calculate electrostatic and van der Waals interaction between atoms within a 14 Å radial distance. The integration timestep was set to 2 fs. Each hydrogen atom and the atom to which it was bonded were similarly constrained and water molecules were made rigid. The timesteps between non-bonded evaluation (nonbondedFreq) were set to 1, specifying how often short-range, non-bonded interactions were calculated. Number of timesteps between each full electrostatic evaluation (fullElectFrequency) was set to 2. Number of steps per cycle was set to 10. Langevin dynamics parameter (langevinDamping) was set to 1 to drive each atom in the system to the target temperature. Periodic boundary conditions were specified. The center of the periodic cell was defined in cellOrigin to which all the coordinates wrapped when wrapAll was set on. Particle Mesh Ewald (PME), applicable only to periodic simulations, was employed as an efficient full electrostatics method that is more accurate and less expensive than larger cutoffs. PME grid dimensions corresponding to the size of the periodic cell were specified. Group-based pressure was used to control the periodic cell fluctuations. The dynamical properties of the barostat and the target pressure were controlled by parameters of the Nosé-Hoover Langevin piston. To initiate the simulation, energy minimization for 5000 steps at constant temperature and pressure was performed for all systems that contained ssDNA, GQD, water molecules, and NaCl ions. After minimization, all systems underwent equilibration for 100 ns.

In all MD simulation figures, GQDs are displayed by line representations and black coloring method. ssDNA secondary structures are displayed by the New Cartoon representation. Adsorbed residue atoms are displayed by the CPK drawing method and red, green, and blue coloring methods for A_30_, C_30_, and T_30_, respectively. Oxidation groups attached to GQDs are represented by the CPK drawing method and magenta coloring method.

## Acknowledgements

We thank Ron Zuckerman and Jae Hong Kim for assistance with AFM imaging and peptoid synthesis. We thank Evan Miller for fluorimeter use, and the Molecular Graphics and Computation Facility at UC Berkeley College of Chemistry (NIH S10OD023532) for access to the computing facility during this work. We acknowledge support of an NIH NIDA CEBRA award # R21DA044010 (to M.P.L.), a Burroughs Wellcome Fund Career Award at the Scientific Interface (CASI) (to M.P.L.), the Simons Foundation (to M.P.L.), a Stanley Fahn PDF Junior Faculty Grant with Award # PF-JFA-1760 (to M.P.L.), a Beckman Foundation Young Investigator Award (to M.P.L.), and a DARPA Young Investigator Award (to M.P.L.). M.P.L. is a Chan Zuckerberg Biohub investigator. R.L.P. acknowledges the support of an NSF Graduate Research Fellowship.

## Author Contributions

S.J., R.L.P., P.P, and M.P.L. conceived the idea and designed experiments. S.J. and R.L.P. performed the experiments. B.D., A.K., D.D., W.Q., and P.P. performed and analyzed molecular dynamics simulations and synthesized high-ox-GQDs. H.S. and M.H. synthesized and characterized no-ox-GQDs. All authors discussed the results and wrote the manuscript.

## References

1. Novoselov, K. S. et al. A roadmap for graphene. Nature 490, 192–200 (2012).

2. Castro Neto, A. H., Guinea, F., Peres, N. M. R., Novoselov, K. S. & Geim, A. K. The electronic properties of graphene. Rev. Mod. Phys. 81, 109–162 (2009).

3. Yoon, H. et al. Intrinsic Photoluminescence Emission from Subdomained Graphene Quantum Dots. Adv. Mater. 28, 5255–5261 (2016).

4. Li, L. & Yan, X. Colloidal Graphene Quantum Dots. J. Phys. Chem. Lett. 1, 2572–2576 (2010).

5. Ponomarenko, L. A. et al. Chaotic Dirac Billiard in Graphene Quantum Dots. Science 320, 356–358 (2008).

6. Fujii, S. & Enoki, T. Cutting of Oxidized Graphene into Nanosized Pieces. J. Am. Chem. Soc. 132, 10034–10041 (2010).

7. Eda, G. et al. Blue Photoluminescence from Chemically Derived Graphene Oxide. Adv. Mater. 22, 505–509 (2010).

8. Shen, J., Zhu, Y., Yang, X. & Li, C. Graphene quantum dots: emergent nanolights for bioimaging, sensors, catalysis and photovoltaic devices. Chem. Commun. 48, 3686–3699 (2012).

9. Sun, H., Wu, L., Wei, W. & Qu, X. Recent advances in graphene quantum dots for sensing. Mater. Today 16, 433–442 (2013).

10. Song, S. H. et al. Highly Efficient Light-Emitting Diode of Graphene Quantum Dots Fabricated from Graphite Intercalation Compounds. Adv. Opt. Mater. 2, 1016–1023 (2014).

11. Zhu, S. et al. Strongly green-photoluminescent graphene quantum dots for bioimaging applications. Chem. Commun. 47, 6858–6860 (2011).

12. Peng, J. et al. Graphene quantum dots derived from carbon fibers. Nano lett. 12, 844–849 (2012).

13. Zheng, M. et al. DNA-assisted dispersion and separation of carbon nanotubes. Nat. Mater. 2, 338–342 (2003).

14. Zhang, J. et al. Molecular recognition using corona phase complexes made of synthetic polymers adsorbed on carbon nanotubes. Nat. Nanotechnol. 8, 959–968 (2013).

15. Zhang, H. et al. Universal fluorescence biosensor platform based on graphene quantum dots and pyrene-functionalized molecular beacons for detection of microRNAs. ACS appl. Mater. Inter. 7, 16152–16156 (2015).

16. Liu, F., Choi, J. Y. & Seo, T. S. Graphene oxide arrays for detecting specific DNA hybridization by fluorescence resonance energy transfer. Biosens. Bioelectron. 25, 2361–2365 (2010).

17. He, S. et al. A Graphene Nanoprobe for Rapid, Sensitive, and Multicolor Fluorescent DNA Analysis. Adv. Funct. Mater. 20, 453–459 (2010).

18. Chang, H., Tang, L., Wang, Y., Jiang, J. & Li, J. Graphene Fluorescence Resonance Energy Transfer Aptasensor for the Thrombin Detection. Anal. Chem. 82, 2341–2346 (2010).

19. Wang, Y., Zhang, L., Liang, R.-P., Bai, J.-M. & Qiu, J.-D. Using Graphene Quantum Dots as Photoluminescent Probes for Protein Kinase Sensing. Anal. Chem. 85, 9148–9155 (2013).

20. Jeon, S.-J., Kwak, S.-Y., Yim, D., Ju, J.-M. & Kim, J.-H. Chemically-Modulated Photoluminescence of Graphene Oxide for Selective Detection of Neurotransmitter by “Turn-On” Response. J. Am. Chem. Soc. 136, 10842–10845 (2014).

21. Wang, Y., Tang, L., Li, Z., Lin, Y. & Li, J. In situ simultaneous monitoring of ATP and GTP using a graphene oxide nanosheet–based sensing platform in living cells. Nat. Protoc. 9, 1944–1955 (2014).

22. Li, X. et al. A “turn-on” fluorescent sensor for detection of Pb2+ based on graphene oxide and G-quadruplex DNA. Phys. Chem. Chem. Phys. 15, 12800–12804 (2013).

23. Wen, Y. et al. A graphene-based fluorescent nanoprobe for silver(I) ions detection by using graphene oxide and a silver-specific oligonucleotide. Chem. Commun. 46, 2596–2598 (2010).

24. Liu, Z., Robinson, J. T., Sun, X. & Dai, H. PEGylated Nanographene Oxide for Delivery of Water-Insoluble Cancer Drugs. J. Am. Chem. Soc. 130, 10876–10877 (2008).

25. Luo, N. et al. PEGylated graphene oxide elicits strong immunological responses despite surface passivation. Nat. Commun. 8, 14537 (2017).

26. Zhang, L., Xia, J., Zhao, Q., Liu, L. & Zhang, Z. Functional Graphene Oxide as a Nanocarrier for Controlled Loading and Targeted Delivery of Mixed Anticancer Drugs. Small 6, 537–544 (2010).

27. Sun, X. et al. Nano-graphene oxide for cellular imaging and drug delivery. Nano Res. 1, 203–212 (2008).

28. Lee, J.-H., Choi, Y.-K., Kim, H.-J., Scheicher, R. H. & Cho, J.-H. Physisorption of DNA Nucleobases on h-BN and Graphene: vdW-Corrected DFT Calculations. J. Phys. Chem. C 117, 13435–13441 (2013).

29. Willems, N. et al. Biomimetic Phospholipid Membrane Organization on Graphene and Graphene Oxide Surfaces: A Molecular Dynamics Simulation Study. ACS Nano 11, 1613–1625 (2017).

30. Li, X., Wang, X., Zhang, L., Lee, S. & Dai, H. Chemically Derived, Ultrasmooth Graphene Nanoribbon Semiconductors. Science 319, 1229–1232 (2008).

31. Wu, M., Kempaiah, R., Huang, P.-J. J., Maheshwari, V. & Liu, J. Adsorption and desorption of DNA on graphene oxide studied by fluorescently labeled oligonucleotides. Langmuir 27, 2731–2738 (2011).

32. Patil, A. J., Vickery, J. L., Scott, T. B. & Mann, S. Aqueous Stabilization and Self-Assembly of Graphene Sheets into Layered Bio-Nanocomposites using DNA. Adv. Mater. 21, 3159–3164 (2009).

33. Varghese, N. et al. Binding of DNA nucleobases and nucleosides with graphene. ChemPhysChem 10, 206–210 (2009).

34. Kim, H. S., Farmer, B. L. & Yingling, Y. G. Effect of Graphene Oxidation Rate on Adsorption of Poly-Thymine Single Stranded DNA. Adv. Mater. Inter. 4, 1601168 (2017).

35. Shi, X., Kong, Y., Zhao, Y. & Gao, H. Molecular dynamics simulation of peeling a DNA molecule on substrate. Acta Mech. Sinica 21, 249–256 (2005).

36. Antony, J. & Grimme, S. Structures and interaction energies of stacked graphene–nucleobase complexes. Phys. Chem. Chem. Phys. 10, 2722–2729 (2008).

37. Lin, Y., Chapman, R. & Stevens, M. M. Integrative Self-Assembly of Graphene Quantum Dots and Biopolymers into a Versatile Biosensing Toolkit. Adv. Funct. Mater. 25, 3183–3192 (2015).

38. Hayakawa, T., Ishii, Y. & Kawasaki, S. Sodium ion battery anode properties of designed graphene-layers synthesized from polycyclic aromatic hydrocarbons. RSC Adv. 6, 22069–22073 (2016).

39. Dong, Y. et al. Blue luminescent graphene quantum dots and graphene oxide prepared by tuning the carbonization degree of citric acid. Carbon 50, 4738–4743 (2012).

40. Kundu, S., Wang, Y., Xia, W. & Muhler, M. Thermal Stability and Reducibility of Oxygen-Containing Functional Groups on Multiwalled Carbon Nanotube Surfaces: A Quantitative High-Resolution XPS and TPD/TPR Study. J. Phys. Chem. C 112, 16869–16878 (2008).

41. Jang, M.-H. et al. Is the chain of oxidation and reduction process reversible in luminescent graphene quantum dots? Small 11, 3773–3781 (2015).

42. LeCroy, G. E. et al. Characteristic Excitation Wavelength Dependence of Fluorescence Emissions in Carbon “Quantum” Dots. J. Phys. Chem. C 121, 28180–28186 (2017).

43. Kruss, S. et al. Neurotransmitter Detection Using Corona Phase Molecular Recognition on Fluorescent Single-Walled Carbon Nanotube Sensors. J. Am. Chem. Soc. 136, 713–724 (2014).

44. Li, M., Zhou, X., Ding, W., Guo, S. & Wu, N. Fluorescent aptamer-functionalized graphene oxide biosensor for label-free detection of mercury(II). Biosens. Bioelectron. 41, 889–893 (2013).

45. Zhao, W., Lin, L. & Hsing, I.-M. Rapid synthesis of DNA-functionalized gold nanoparticles in salt solution using mononucleotide-mediated conjugation. Bioconjugate Chem. 20, 1218–1222 (2009).

46. Zheng, S., Tu, Q., Urban, J. J., Li, S. & Mi, B. Swelling of Graphene Oxide Membranes in Aqueous Solution: Characterization of Interlayer Spacing and Insight into Water Transport Mechanisms. ACS Nano 11, 6440–6450 (2017).

47. Lu, C., Huang, P.-J. J., Liu, B., Ying, Y. & Liu, J. Comparison of Graphene Oxide and Reduced Graphene Oxide for DNA Adsorption and Sensing. Langmuir 32, 10776–10783 (2016).

48. Sowerby, S. J., Cohn, C. A., Heckl, W. M. & Holm, N. G. Differential adsorption of nucleic acid bases: Relevance to the origin of life. Proc. Natl. Acad. Sci. 98, 820–822 (2001).

49. Manohar, S. et al. Peeling Single-Stranded DNA from Graphite Surface to Determine Oligonucleotide Binding Energy by Force Spectroscopy. Nano Lett. 8, 4365–4372 (2008).

50. Huang, Z. & Liu, J. Length-Dependent Diblock DNA with Poly-cytosine (Poly-C) as High-Affinity Anchors on Graphene Oxide. Langmuir 34, 1171–1177 (2018).

51. Yang, D. et al. Chemical analysis of graphene oxide films after heat and chemical treatments by X-ray photoelectron and Micro-Raman spectroscopy. Carbon 47, 145–152 (2009).

52. Zuckermann, R. N., Kerr, J. M., Kent, S. B. H. & Moos, W. H. Efficient method for the preparation of peptoids [oligo(N-substituted glycines)] by submonomer solid-phase synthesis. J. Am. Chem. Soc. 114, 10646–10647 (1992).

53. Nam, K. T. et al. Free-floating ultrathin two-dimensional crystals from sequence-specific peptoid polymers. Nat. Mater. 9, 454 (2010).

54. Chio, L. et al. Electrostatic-assemblies of single-walled carbon nanotubes and sequence-tunable peptoid polymers detect a lectin protein and its target sugars. Nano Lett. (2019). doi:10.1021/acs.nanolett.8b04955

55. Phillips, J. C. et al. Scalable molecular dynamics with NAMD. J. Comput. Chem. 26, 1781–1802 (2005).

56. Kumar, S. et al. Scalable molecular dynamics with NAMD on the IBM Blue Gene/L system. IBM J. Res. Dev. 52, 177–188 (2008).

57. Brooks, B. R. et al. CHARMM: A program for macromolecular energy, minimization, and dynamics calculations. J. Comput. Chem. 4, 187–217 (1983).

58. Humphrey, W., Dalke, A. & Schulten, K. VMD: Visual molecular dynamics. J. Mol. Graph. 14, 33–38 (1996).

59. van Dijk, M. & Bonvin, A. M. J. J. 3D-DART: a DNA structure modelling server. Nucleic Acids Res. 37, W235–W239 (2009).

60. Pedretti, A., Villa, L. & Vistoli, G. VEGA – An open platform to develop chemo-bio-informatics applications, using plug-in architecture and script programming. J. Comput. Aided Mol. Des. 18, 167–173 (2004).

61. Jorgensen, W. L., Chandrasekhar, J., Madura, J. D., Impey, R. W. & Klein, M. L. Comparison of simple potential functions for simulating liquid water. J. Chem. Phys. 79, 926–935 (1983).

