## Supplemental Information for "Graphene Quantum Dot Oxidation Governs Noncovalent Biopolymer Adsorption"

‡These authors contributed equally

### Supplementary Figures

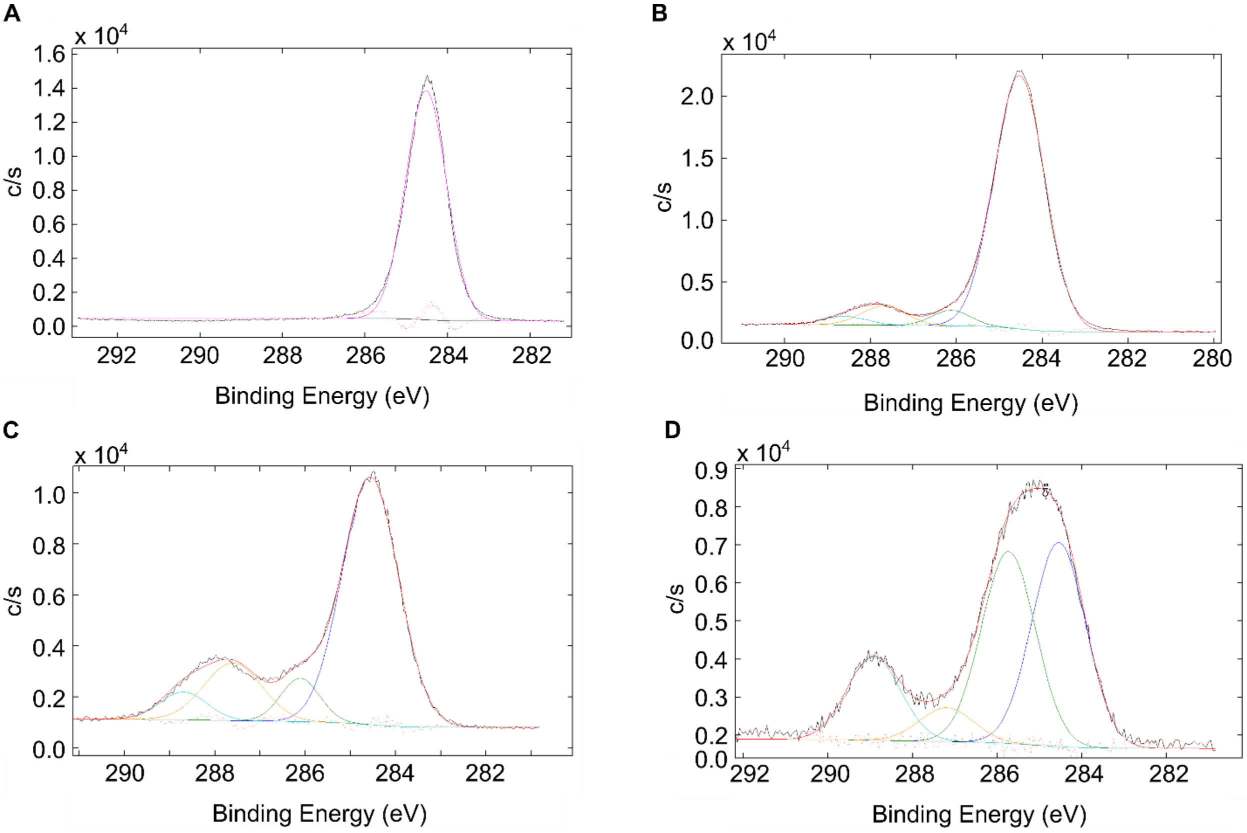


**E**

| Sample | C-C (284.5 eV) | C-O (286.1 eV) | C=O (287.5 eV) | O-C=O (288.7 eV) |
|---|---|---|---|---|
| No-ox-GQD | 100 | 0 | 0 | 0 |
| Low-ox-GQD | 87.55 | 4.06 | 5.84 | 2.56 |
| Med-ox-GQD | 68.95 | 8.31 | 16.36 | 6.38 |
| High-ox-GQD | 39.78 | 37.4 | 6.57 | 16.24 |

**Figure S1**. **Deconvoluted carbon 1s (C1s) X-ray photoelectron spectroscopy (XPS) characterization of GQDs. (A)** no-ox-GQD, **(B)** low-ox-GQD, **(C)** med-ox-GQD, and **(D)** high-ox-GQD samples included in Fig. 1B. **(E)** Relative peak areas for C1s chemical bonds in each sample.

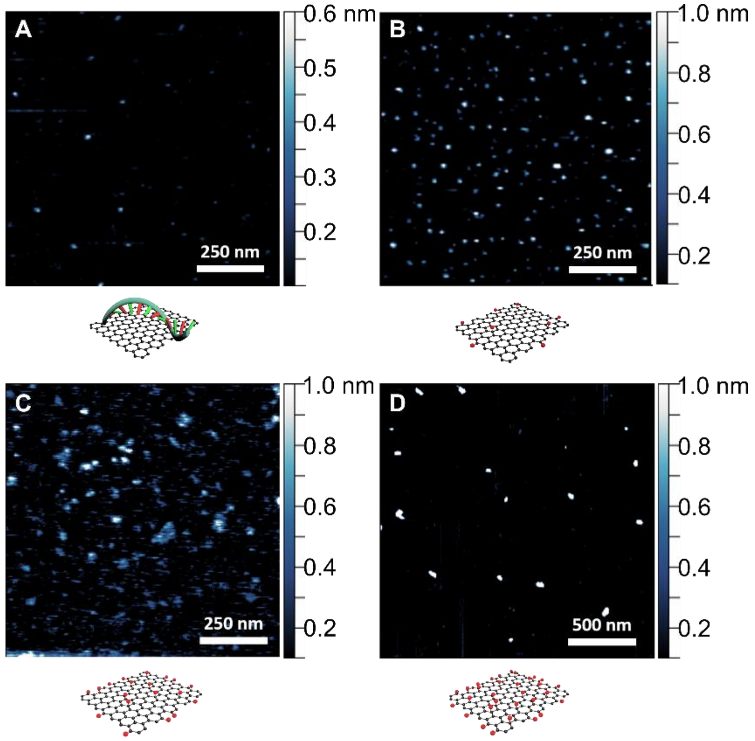


**Figure S2**. **AFM characterization of GQDs.** AFM images and accompanying schematics for **(A)** $(GT)_{15}$-no-ox-GQDs, **(B)** low-ox-GQDs, **(C)** med-ox-GQDs, and **(D)** high-ox GQDs.

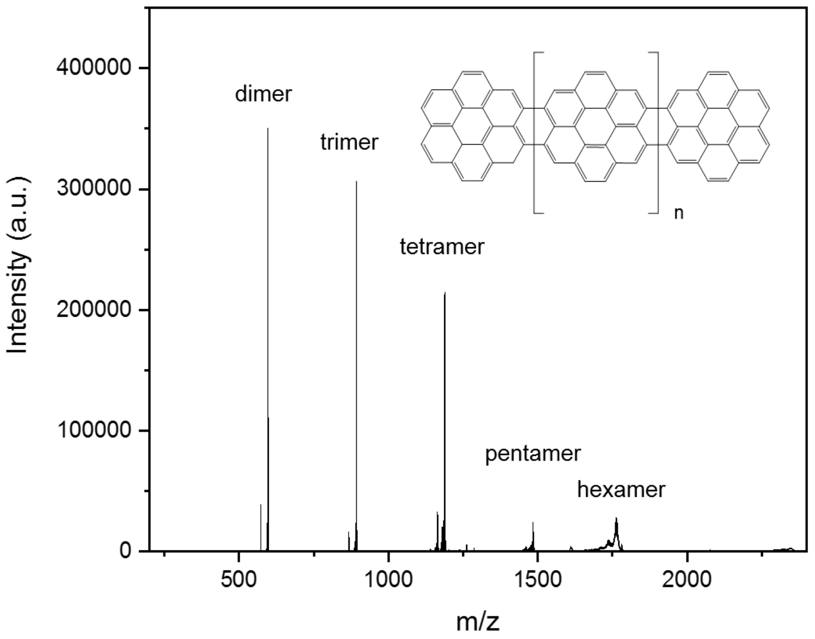


**Figure S3**. **Matrix-assisted laser desorption/ionization time-of-flight mass spectroscopy (MALDI-TOF MS) characterization of no-ox-GQDs**. Peak bands observed at m/z = 596, 892, 1188, 1484, and 1780 are attributed to planar coronene dimer, trimer, tetramer, and pentamer, and hexamer structures, respectively.

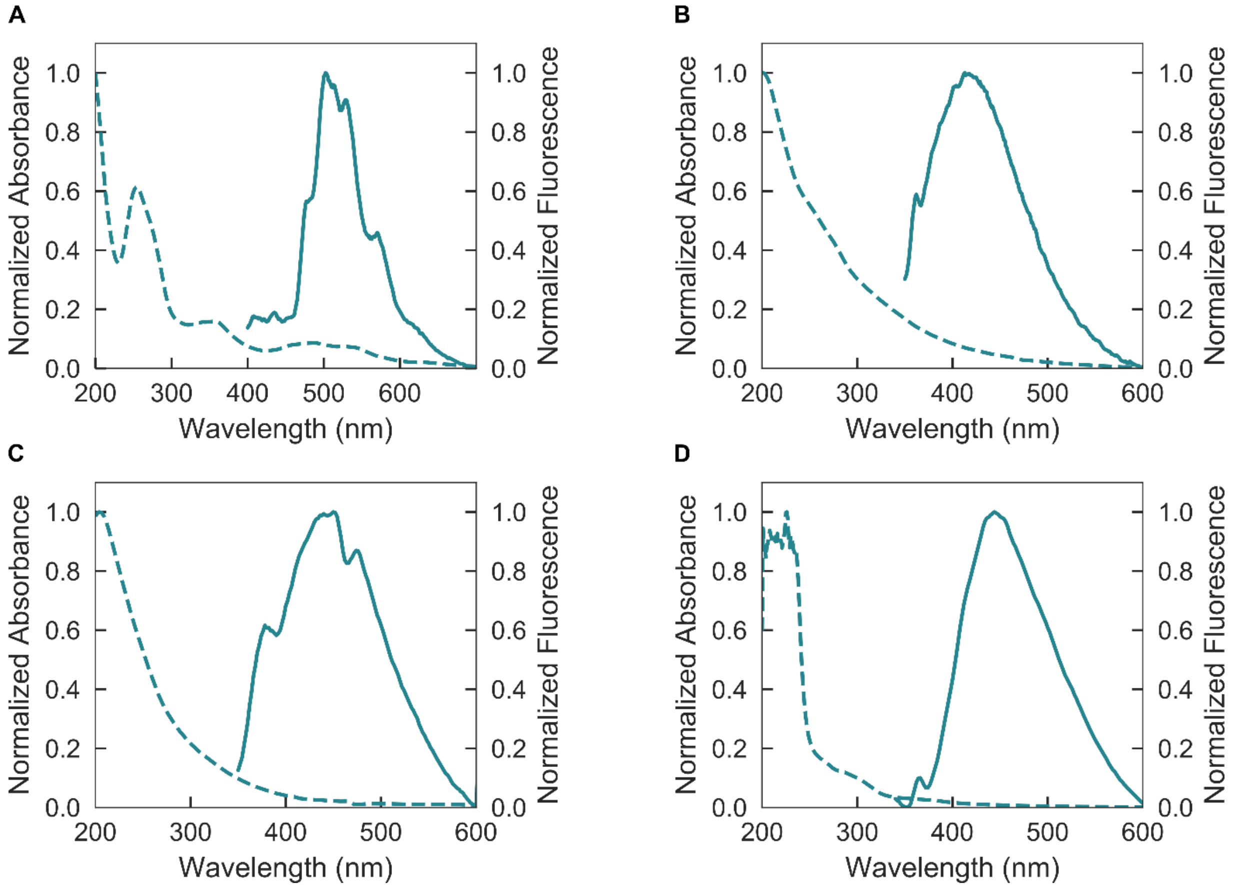


**Figure S4**. **Optical absorption and emission characterization of GQDs.** Normalized absorption (dashed) and fluorescence emission (solid) spectra of **(A)** $(GT)_{15}$-no-ox-GQDs (excitation 340 nm), **(B)** low-ox-GQDs (excitation 320 nm), **(C)** med-ox-GQDs (excitation 320 nm), and **(D)** high-ox-GQDs (excitation 340 nm) in water.

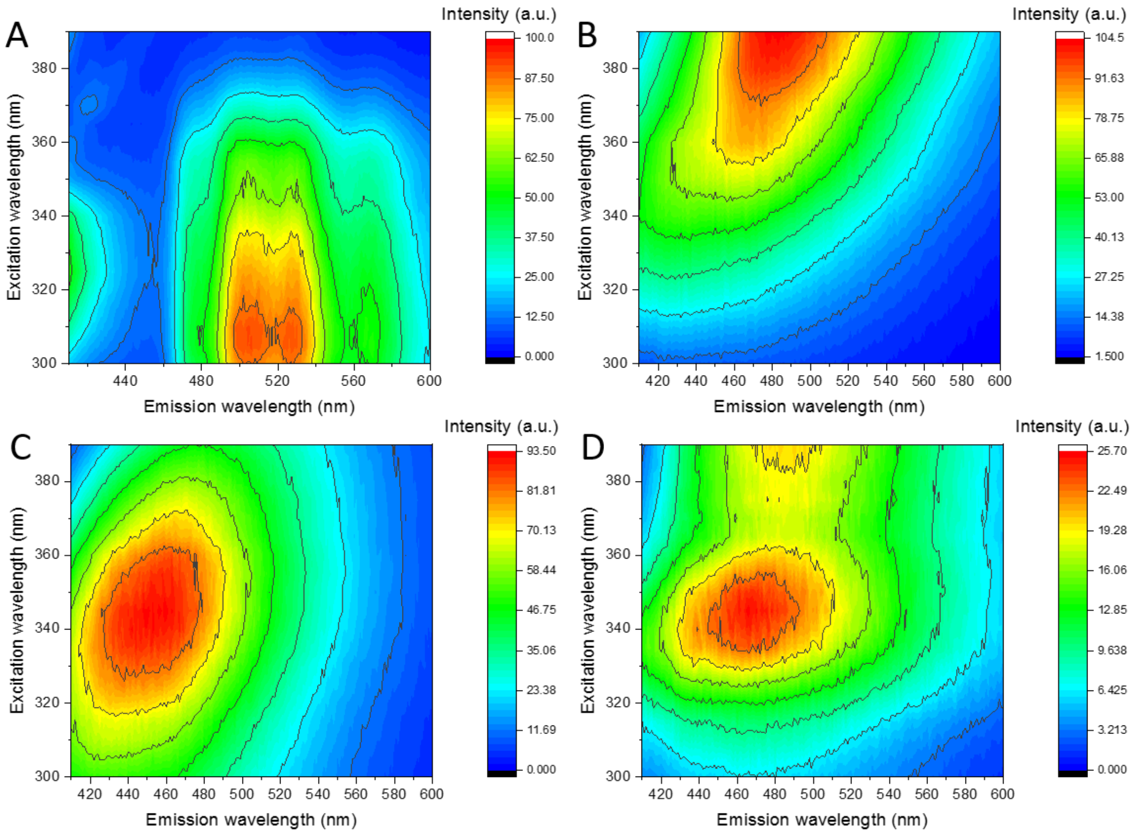


**Figure S5. Excitation-emission profiles of GQDs. (A)** $(GT)_{15}$-no-ox-GQDs, **(B)** low-ox-GQDs, **(C)** med-ox-GQDs, and **(D)** high-ox-GQDs in water. All spectra were collected in intervals of 5 nm for excitation wavelength.

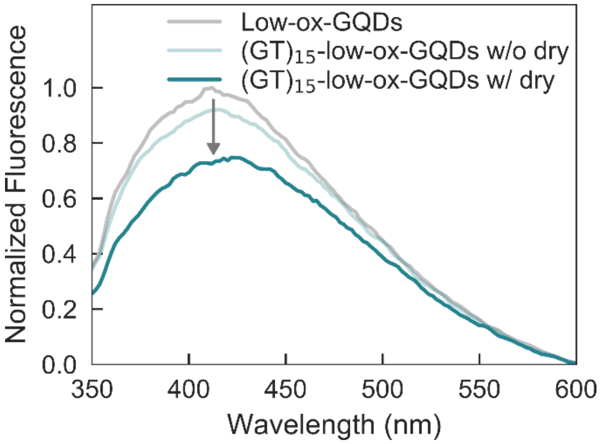


**Figure S6**. **Effectiveness of vacuum drying for ssDNA adsorption to low-ox-GQDs.** Normalized fluorescence emission spectra of low-ox-GQDs after vacuum evaporation (gray), $(GT)_{15}$ and low-ox-GQD mixture without vacuum evaporation (light blue), and $(GT)_{15}$ and low-ox-GQD mixture with vacuum evaporation (dark blue). Concentrations of all low-ox-GQD samples are equivalent.

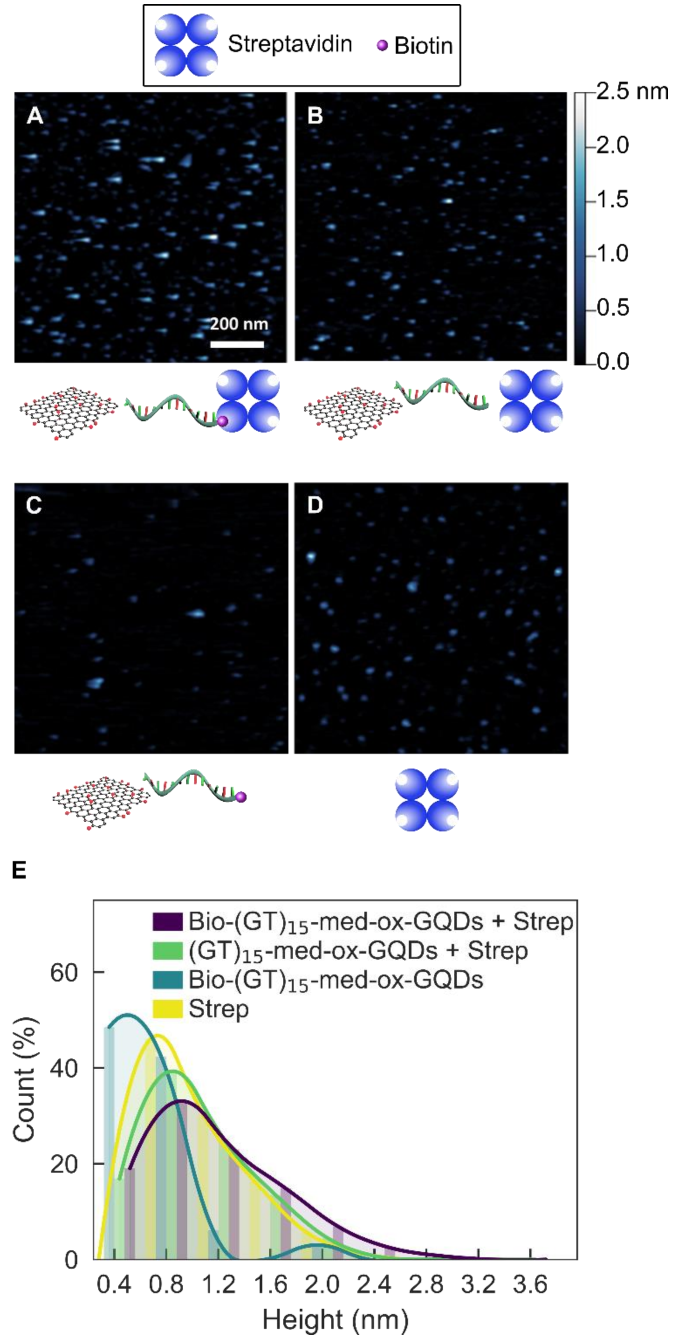


**Figure S7. AFM verification of no ssDNA adsorption on med-ox-GQDs.** AFM images and accompanying schematics for **(A)** biotinylated-$(GT)_{15}$-med-ox-GQDs and streptavidin (Bio-$(GT)_{15}$-med-ox-GQD + Strep), **(B)** $(GT)_{15}$-med-ox-GQDs and streptavidin ($(GT)_{15}$-med-ox-GQD + Strep), **(C)** biotinylated-$(GT)_{15}$-med-ox-GQDs (Bio-$(GT)_{15}$-med-ox-GQD), and **(D)** streptavidin (Strep). **(E)** Corresponding height distribution histograms. Bin width is 0.4 nm and curve fits are added to guide the eye.

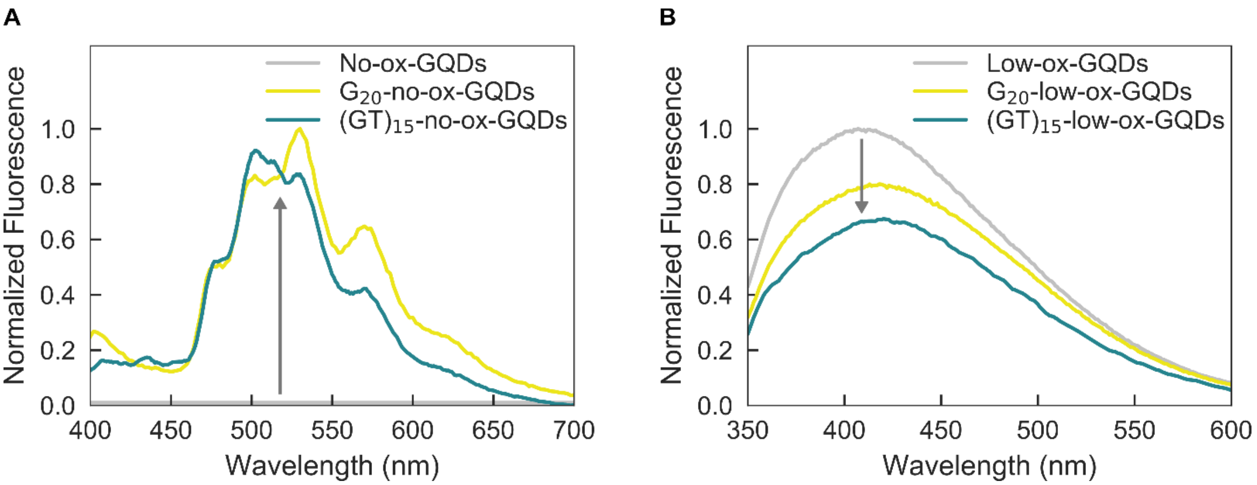


**Figure S8. Normalized fluorescence emission spectra to probe ssDNA sequence dependence for adsorption to GQDs.** Fluorescence of $G_{20}$-GQDs (yellow) compared to $(GT)_{15}$-GQDs (blue) for **(A)** no-ox-GQDs and **(B)** low-ox-GQDs. All GQD fluorescence spectra are normalized by the absorbance at 320 nm.

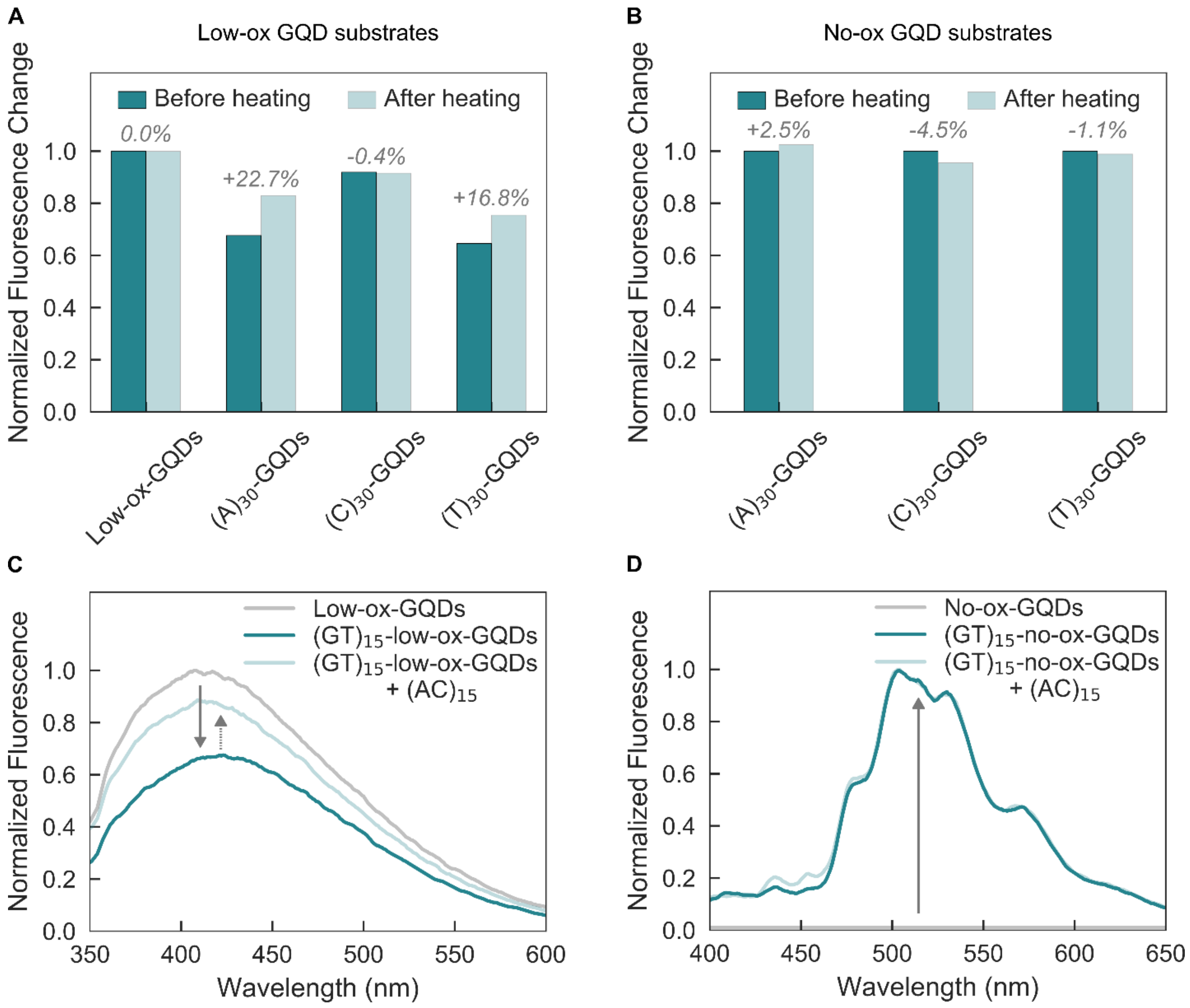


**Figure S9**. **ssDNA desorption from low- and no-ox-GQDs indicates strength of noncovalent binding interactions is inversely proportional to GQD oxidation level.** Fluorescence intensity change of **(A)** ssDNA-low-ox-GQDs and **(B)** ssDNA-no-ox-GQDs induced by thermal desorption of ssDNA after heating ssDNA-GQD samples at 50 ℃ for 2 hours. Normalized fluorescence emission spectra of **(C)** $(GT)_{15}$-low-ox-GQDs and **(D)** $(GT)_{15}$-no-ox-GQDs, after adding five-fold excess of complementary ssDNA, $(AC)_{15}$. All GQD fluorescence spectra are normalized by the absorbance at 320 nm.

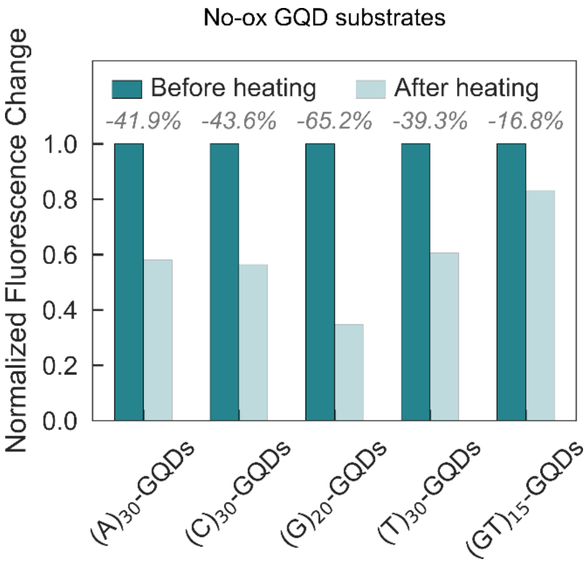


**Figure S10. Thermally-induced ssDNA desorption from ssDNA-no-ox-GQDs.** Fluorescence intensity change of ssDNA-no-ox-GQDs induced by thermal desorption of ssDNA after heating the mixture at 95 ℃ for 2 hours.

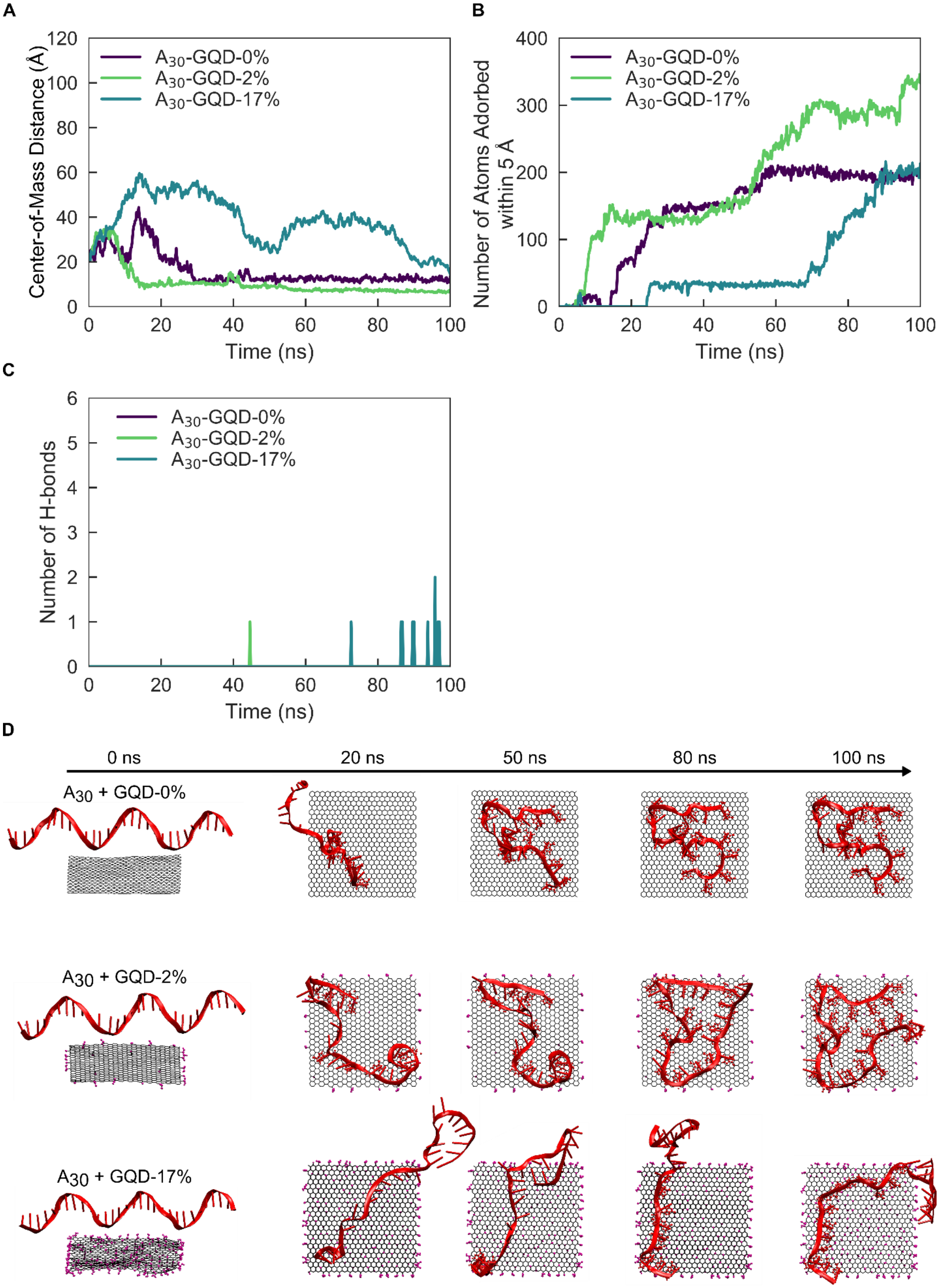


**Figure S11. Molecular dynamics simulations of $A_{30}$ ssDNA adsorbing to GQDs of varying oxidation levels.** Time-dependent **(A)** center-of-mass distance, **(B)** number of atoms adsorbed within 5 Å of the GQD surface, and **(C)** number of hydrogen bonds for $A_{30}$ ssDNA adsorbing to GQD-0%, GQD-2%, and GQD-17%. **(D)** Initial (left) and final (right) configurations of $A_{30}$ ssDNA with GQD-0%, GQD-2%, and GQD-17% for a 100 ns simulation.

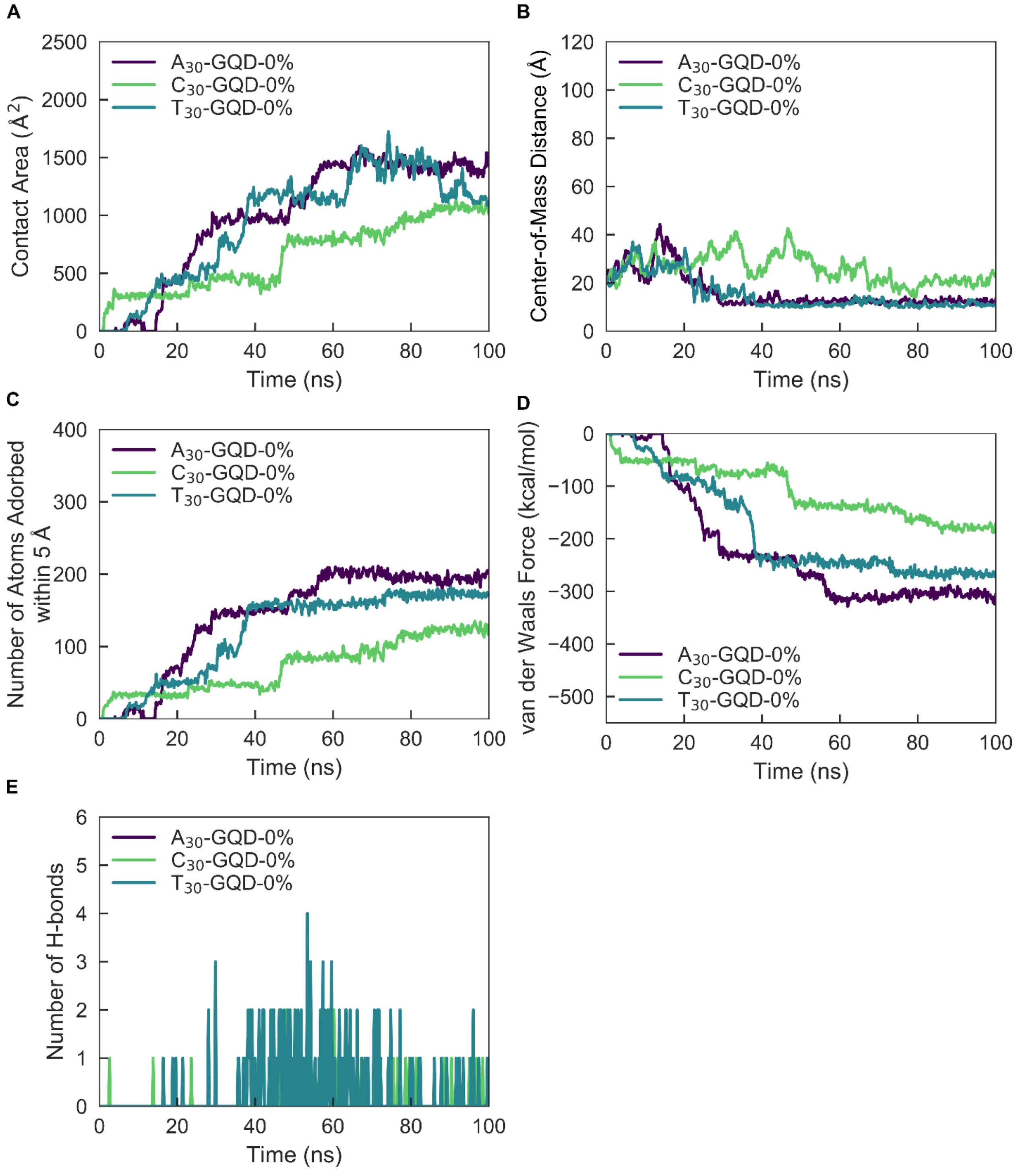
A
B
C
D
E
Contact Area (Å²)
Center-of-Mass Distance (Å)
Number of Atoms Adorbed within 5 Å
van der Waals Force (kcal/mol)
Number of H-bonds
Time (ns)
A30-GQD-0%
C30-GQD-0%
T30-GQD-0%

**F**

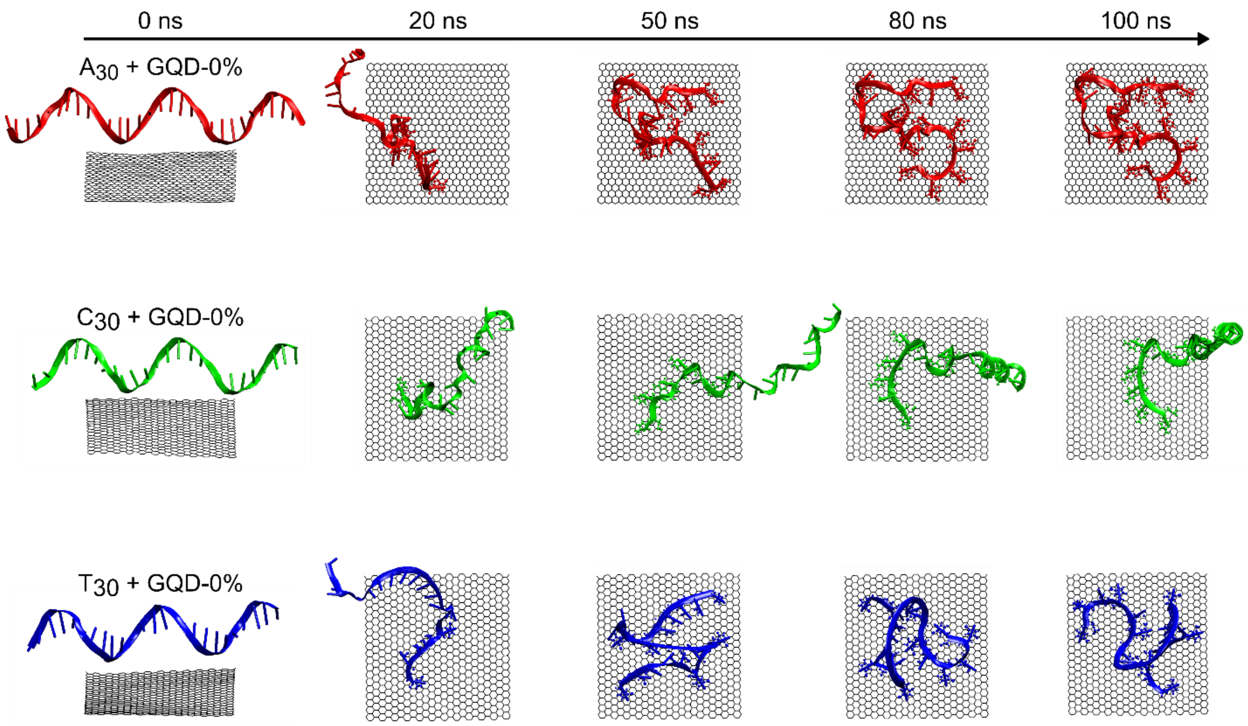


**Figure S12. Molecular dynamics simulations of $A_{30}$, $C_{30}$, and $T_{30}$ ssDNA adsorbing to GQDs without oxidation, GQD-0%.** Time-dependent **(A)** contact area, **(B)** center-of-mass distance, **(C)** number of atoms adsorbed within 5 Å of the GQD surface, **(D)** van der Waals interactions, and **(E)** number of hydrogen bonds for $A_{30}$, $C_{30}$, and $T_{30}$ ssDNA adsorbing to GQD-0%. **(F)** Initial (left) and final (right) configurations of ssDNA with GQD-0% for a 100 ns simulation.

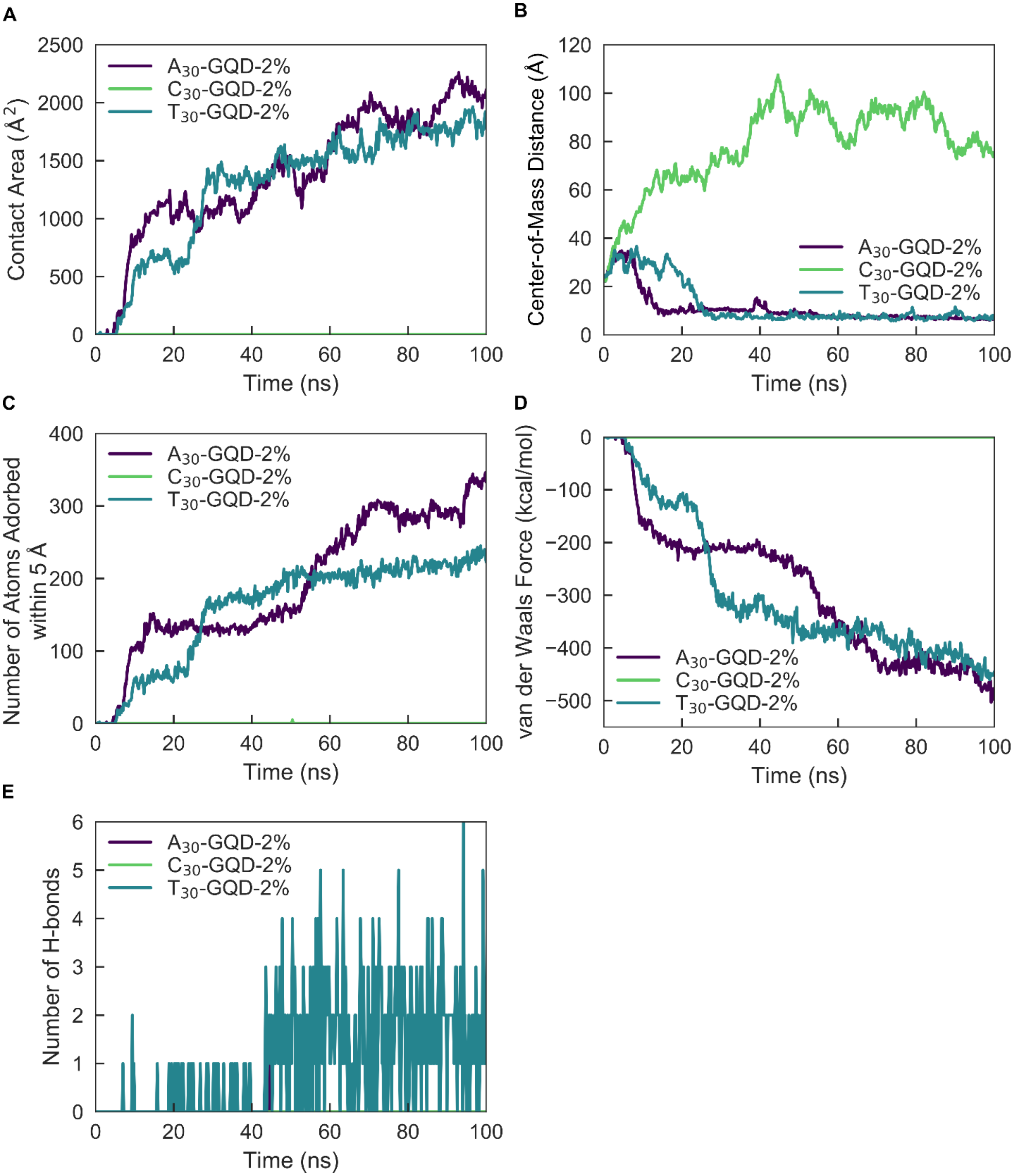
A
B
C
D
E
Contact Area (Å$^2$)
Center-of-Mass Distance (Å)
Number of Atoms Adorbed within 5 Å
van der Waals Force (kcal/mol)
Number of H-bonds
Time (ns)
A30-GQD-2%
C30-GQD-2%
T30-GQD-2%

F

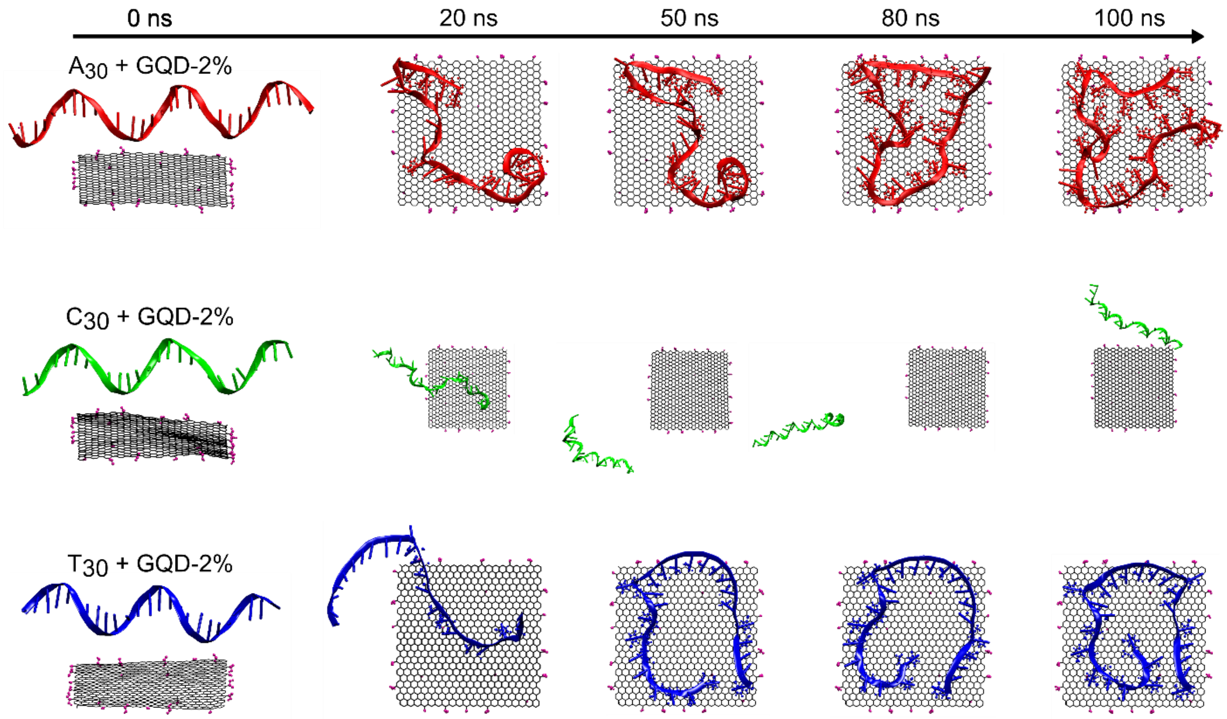


**Figure S13. Molecular dynamics simulations of $A_{30}$, $C_{30}$, and $T_{30}$ ssDNA adsorbing to GQDs low oxidation, GQD-2%.** Time-dependent **(A)** contact area, **(B)** center-of-mass distance, **(C)** number of atoms adsorbed within 5 Å of the GQD surface, **(D)** van der Waals interactions, and **(E)** number of hydrogen bonds for $A_{30}$, $C_{30}$, and $T_{30}$ ssDNA adsorbing to GQD-2%. **(F)** Initial (left) and final (right) configurations of ssDNA with GQD-2% for a 100 ns simulation. Note zoomed-out view of $C_{30}$ due to larger distance of ssDNA to GQD surface; GQD size is the same throughout.
